# The Ubiquitin–Proteasome System is an Important Driver of EBV-Associated Nasopharyngeal Carcinoma Progression: A Meta-Analysis of Transcriptomics Data

**DOI:** 10.1101/2025.08.30.673297

**Authors:** Hana Ratnawati, Ardo Sanjaya, Aldrich Christiandy, Lawrence S Young, Sascha Ott

## Abstract

**Background:** Epstein–Barr virus (EBV)-associated nasopharyngeal carcinoma (NPC) is characterized by extensive immune infiltration, yet immune evasion remains a hallmark of the disease. In this study, we aimed to leverage publicly available datasets to identify EBV–host gene interactions and re-map their expression at single-cell resolution.

**Methods:** We conducted a meta-analysis of transcriptomic datasets to identify differentially expressed genes in NPC, mapped these genes to EBV–host interaction data, and constructed a network. Network clustering and pathway enrichment were performed, and single-cell RNA sequencing datasets were used to map these genes at cellular resolution. We analysed differences in cell cycle, immune signaling, cell–cell interactions, and assessed prognostic associations of a 12-gene UPS signature using the GEPIA2 platform.

**Results:** We identified 85 EBV-interacting DEGs, most regulated by lytic-phase proteins. MCODE clustering highlighted genes related to the ubiquitin–proteasome system (UPS). Single-cell analyses confirmed elevated UPS-related gene expression in NPC compared to EBV-negative oropharyngeal cancer. UPS-High cells exhibited lower proliferative activity, enriched stemness signaling, and reduced antigen presentation and immune activation, whereas UPS-Low cells showed marked upregulation of growth-promoting pathways. Survival analysis revealed that high UPS expression was associated with poorer outcomes in both pan-cancer and head and neck squamous cell carcinoma datasets. The proportion of UPS-High cells varied widely across patients.

**Conclusion:** UPS activity, influenced by EBV lytic proteins, is linked to immune evasion, stemness, proliferation, and adverse prognosis. These findings support UPS as a potential biomarker and therapeutic target in EBV-associated NPC, warranting NPC-specific validation and functional studies.

**Graphical Abstract:** 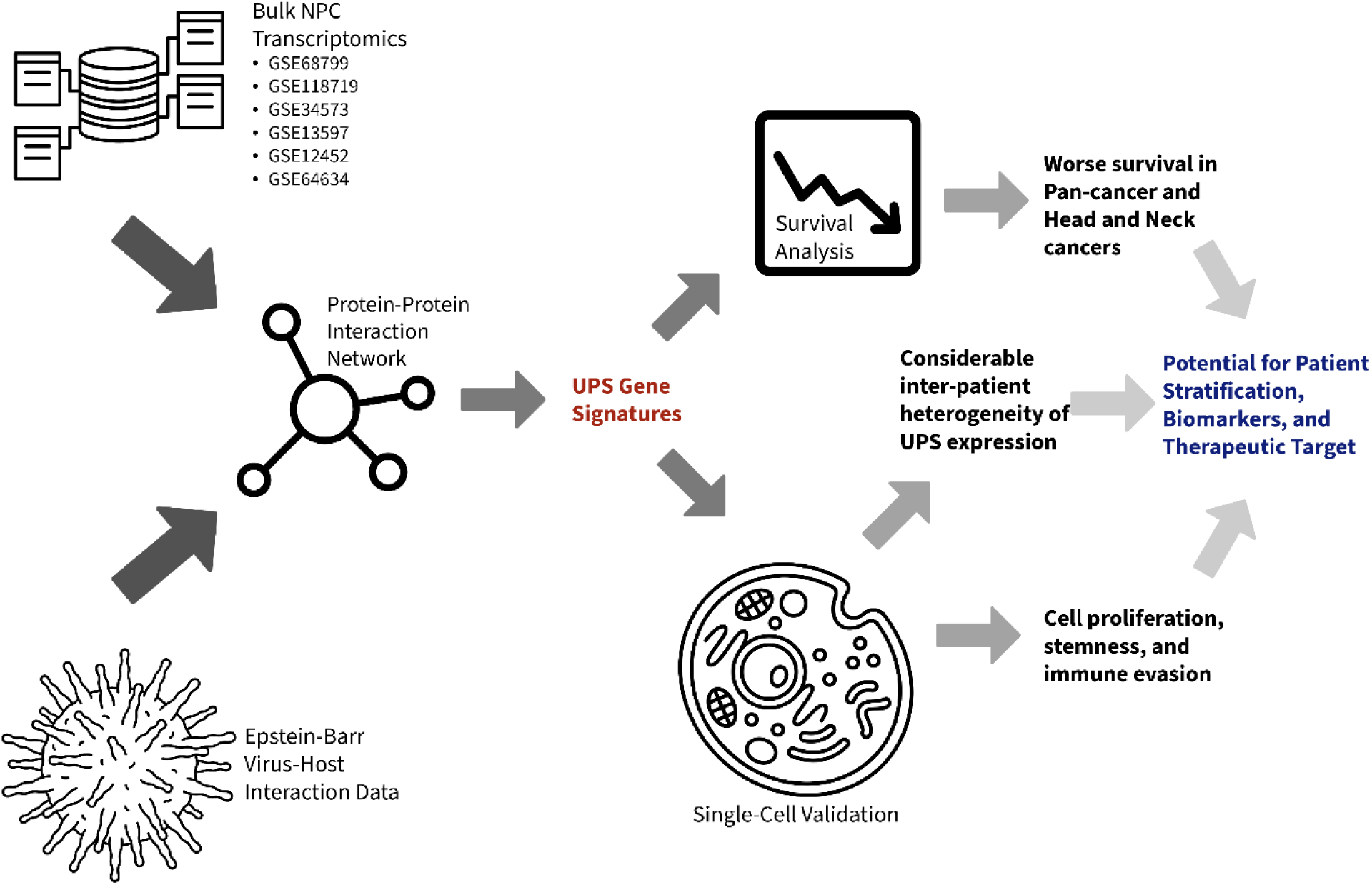

EBV-driven ubiquitin–proteasome dysregulation in nasopharyngeal carcinoma revealed by bulk and single-cell transcriptomics, linked to immune evasion and poor survival.

## Introduction

Nasopharyngeal carcinoma (NPC) is a relatively rare but highly aggressive malignancy arising from the epithelial lining of the nasopharynx. In Southeast Asia and China, it ranks among the top ten cancers by incidence and mortality, with age-standardized incidence rates in China of 2.4 per 100,000 person-year, and in Indonesia, 6.2 per 100,000. However, the age-standardized mortality rate due to NPC in Indonesia is the highest in Southeast Asian countries, at 4.3 per 100,000, triple the mortality rate in China. An estimated 87,000 new cases and 51,000 deaths occur annually, underscoring its significant regional burden [1–3]. According to the WHO, an estimated 120,434 new NPC cases and 73,482 deaths occurred in 2022. Therefore, A better understanding of the molecular mechanisms underlying NPC is critical to improving diagnostics and developing new therapeutic targets.

A defining feature of NPC in endemic regions is its consistent association with Epstein–Barr virus (EBV) infection, especially in the non-keratinizing subtype. EBV is a gamma-herpesvirus capable of latent and lytic infection in epithelial cells. Although the lytic phase typically triggers full viral replication and immune activation, it is transient in NPC and may still contribute to genomic instability and inflammation. [4–6] In contrast, latent infection predominates in NPC, characterized by restricted viral gene expression, including EBNA1, LMP1, LMP2, EBERs, and BART microRNAs, which modulate oncogenesis and immune evasion [2,7]

A hallmark of EBV-associated NPC is the dense infiltration of immune cells, such as neutrophils, natural killer (NK) cells, regulatory T cells (Tregs), macrophages, dendritic cells, and B cells. [8] Although tumor-infiltrating leukocytes (TILs) are generally linked to improved outcomes in many cancers, their role in NPC is more complex. Despite the abundant immune presence, these cells often fail to mount effective anti-tumor responses, likely due to EBV-mediated immunosuppression. [9] The virus manipulates the TME through various immunomodulatory mechanisms. EBV induces immunosuppressive cytokines such as IL-10, TGF-β, and CCL20, which impair cytotoxic T cell activity and encourage regulatory immune subsets. [10,11] Among these latent genes, LMP1 (Latent Membrane Protein 1) plays a pivotal role by activating key survival and immune-modulating pathways such as NF-κB, JAK/STAT, and PI3K/AKT, driving tumor progression. [12,13] LMP1 also upregulates PD-L1 expression, fostering T cell exhaustion. [14] LMP2A, another latent protein, activates the PI3K/AKT and MAPK pathways and promotes epithelial–mesenchymal transition (EMT), enhancing invasiveness. [15,16] EBV non-coding RNAs, such as EBERs, activate IL-6 and IL-10 production via pattern recognition receptors like RIG-I and TLR3, while also dampening type I interferon responses. [17,18] EBV-encoded BART microRNAs also suppress immune activation by targeting host transcripts involved in antigen processing and interferon signaling. [19] Together, these EBV-induced alterations profoundly impact the immune-dense tumor microenvironment (TME) of NPC, transforming the TME into an immunosuppressive niche that facilitates immune evasion, tumor survival, and metastasis. This paradox makes EBV-associated NPC a valuable model for exploring immune escape mechanisms in cancer.

Another pathway by which EBV supports oncogenesis is through manipulation of the ubiquitin– proteasome system (UPS), a key regulator of protein degradation, immune surveillance, apoptosis, and cell cycle control. LMP1 itself is regulated by the UPS, but this protein also regulates the ubiquitination of several proteins, such as TRAF6, NF-κB, and CHIP, resulting in its alteration of function. [20] The protein LMP1 is also involved in regulating many proteins, including LUBAC, LIMD1, and p62, which mediate selective autophagy and also disturb the ubiquitination process. [21]The UPS is central to antigen presentation as many viral antigens are processed and presented via MHC class I molecules. [22,23] Furthermore, EBV interferes with host tumor suppressor mechanisms via UPS modulation. [22] The lytic protein BPLF1 has deubiquitinase activity that suppresses pathways, including RIG-I and cGAS, thereby impairing interferon responses. [23] Altogether, EBV not only initiates NPC pathogenesis but also sustains tumor progression by actively reshaping immune and stromal dynamics. Its complex interactions with host immunity, signaling, and protein regulation suggest that targeting EBV-specific pathways, such as viral latency genes, immune checkpoints, or UPS interference, could offer novel therapeutic strategies.

To explore these mechanisms at scale, transcriptomic analysis provides a valuable approach. We aim to leverage bulk RNA sequencing data integrated through a meta-analytic approach to allow robust identification of differentially expressed genes (DEGs) across multiple datasets. These DEGs will subsequently be filtered to identify genes encoding proteins that interact with EBV-expressed proteins. The identified genes will then be mapped onto single-cell RNA sequencing data to investigate their expression at a cellular resolution. This approach is used to help elucidate the cell types in which these genes are most active and provide insights into the interplay between EBV infection and the tumor microenvironment. This integrative strategy allows us to better understand the role of EBV in NPC pathogenesis and identify novel therapeutic candidates in the context of viral oncogenesis.

## Methods

### RNA-Seq and Microarray datasets

Publicly available RNA-Seq and microarray datasets were obtained from the GEO database (https://www.ncbi.nlm.nih.gov/geo) using the following accession numbers: GSE68799, GSE118719 [24], GSE34573 [25], GSE13597 [26], GSE12452 [27], and GSE64634 [28] (Table 1). Each dataset was individually analyzed for differentially expressed genes (DEGs) using the limma (RRID:SCR_010943) R package [10], comparing normal vs cancerous tissue. P-values were adjusted using the false discovery rate (FDR) correction method to account for multiple hypothesis tests. Significant DEGs from RNA-Seq and microarray experiments were subsequently grouped for downstream meta-analysis.

**Table 1.** Publicly available transcriptomics datasets used in the meta-analysis.

| GEO Accession Codes | Experiment Type | Experimental Design |
| --- | --- | --- |
| **GSE68799** | RNA-Seq | 42 nasopharyngeal carcinomas;<br>4 normal mucosae |
| **GSE118719** | RNA-Seq | 7 nasopharyngeal carcinomas;<br>4 normal mucosae |
| **GSE34573** | Microarray | 16 nasopharyngeal carcinomas;<br>4 normal controls |
| **GSE13597** | Microarray | 25 nasopharyngeal carcinomas;<br>3 normal controls |
| **GSE12452** | Microarray | 31 nasopharyngeal carcinomas;<br>10 normal controls |
| **GSE64634** | Microarray | 12 nasopharyngeal carcinomas;<br>10 normal controls |

### Meta-regression analysis of DEGs

To account for the heterogeneity of datasets from RNA-Seq and microarray procedures, a meta-regression was conducted on the DEGs using the logFC as the measure of effect size and the standard error as the measure of variance. The standard errors were Winsorized at the 5^th^ and 95^th^ percentiles to minimize extreme values. The meta-regression was conducted using a random-effects model with the restricted maximum likelihood (REML) estimation method, incorporating platform type as a moderator to account for batch effects. For each gene, the following model was applied:

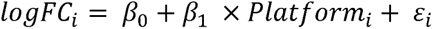

β_0_ represents the estimated overall effect size, β_1_ represents the platform-specific variations, and ε_i_ represents residual variance of platform *i*. The model was implemented with robust variance estimation to handle within-study correlations. For genes where the model failed to converge, effect sizes and p-values were set to NA. Using the Benjamini-Hochberg method, a false discovery rate correction was applied to the p-values. All analyses were conducted in R using the metafor package. (Viechtbauer, 2010).

### EBV Virus-Host Protein-Protein Interaction Network Analysis

Virus-host protein-protein interaction (PPI) data were obtained from the VirHostNet (RRID: SCR_005978) database (https://virhostnet.prabi.fr/) to identify host proteins interacting with EBV proteins. The PPI dataset overlapped with the DEGs previously identified from the meta-regression analysis. The resulting overlaps were then visualized using Cytoscape (RRID: SCR_003032) [31] and processed using the Molecular Complex Detection (MCODE) algorithm (RRID: SCR_015828) [32] to identify the highest interconnected clusters. Identified clusters were enriched using the Reactome database (Milacic et al., 2024) to elucidate key biological processes and pathways associated with EBV-host interactions.

### Single-Cell RNA Sequencing Analysis

Single-cell RNA sequencing (scRNA-seq) datasets were obtained from the GEO database using the following accession numbers: GSE150825, [34] GSE162025, [35] GSE150430, [36] GSE226620, [37] and GSE182227. [38] The datasets GSE150825, GSE162025, and GSE150430 represent nasopharyngeal carcinoma (NPC), while GSE226620 and GSE182227 represent oropharyngeal carcinoma (OPC). OPC datasets were included as EBV-negative cancer controls due to their similar anatomical location and comparable typical histological characteristics to NPC. Enabling direct comparison between EBV-associated NPC and EBV-negative head and neck cancers. Each dataset was processed using Seurat (RRID: SCR_016341) [39] in R through normalization, scaling, and dimensional reduction measures such as principal component analysis (PCA) and Uniform Manifold Approximation and Projection (UMAP). The dataset GSE150430 was pre-normalized and thus directly used in downstream analysis. The datasets were integrated using the Harmony algorithm [40] to correct batch effects and differences in pathological conditions. After integration, the cells were clustered using the FindClusters function, and marker genes were identified using FindAllMarkers using Seurat. The top three marker genes were selected for further enrichment to identify cluster identities. The enrichR (RRID: SCR_001575) [41] package was used to enrich the top marker genes from each cluster against multiple reference databases. Two independent authors verified the cluster identities to ensure accuracy.

### Copy Number Variation Analysis using InferCNV

The InferCNV (RRID: SCR_021140) R package [42] assessed large-scale chromosomal alterations in single cells. This analysis aimed to identify genomic instability in the cell clusters. The following immune and stromal cells were used as reference groups: endothelial, dendritic, and regulatory T cells. The cells with heavy genomic instability, as shown by increased or decreased predicted copy number variations, were then classified as cancer cells.

### Immune Pathway Correlation Analysis

To evaluate pathway activity at the single-cell level, scoring was performed using the AUCell (RRID: SCR_021327) package (v1.20.1) with gene sets from the Gene Ontology Biological Process from the MSigDB (RRID: SCR_016863) database. Rankings of gene expression were computed using AUCell. The top 100 most variable gene sets were selected, and their AUC scores were correlated with the average expression of the UPS gene signature (CCT6A, CCT3, PSMA1, PSMC4, PSMA7, PSMA6, PSMA4, PSMD1, PSMD14, PSMD12, UCHL5, and CCT5) using Spearman’s correlation. For each gene set, both the correlation coefficient (ρ) and the corresponding p-value were computed, and the p-values were adjusted using the Benjamini-Hochberg method. To highlight immunologically relevant biological processes, GO terms containing immune-related keywords (e.g., “IMMUNE”, “T CELL”, “CYTOKINE”, etc.) were filtered from the correlation results. The top 15 positively and top 15 negatively correlated immune-related gene sets were visualized.

### Cancer Cell Reclustering and Classification

Cells were reclustered to investigate the role of proteasomal function in nasopharyngeal carcinoma, and the expression of genes associated with the ubiquitin-proteasome system (UPS) was measured. Specifically, the average expression of CCT6A, CCT3, PSMA1, PSMC4, PSMA7, PSMA6, PSMA4, PSMD1, PSMD14, PSMD12, UCHL5, and CCT5 was calculated, and the cells were classified into UPS-high and UPS-low groups based on the expression of these proteasomal genes. The cell groups were further assessed for cell proliferation by calculating the cell cycle states in Seurat. [39] The proportion of cells in G1, G2/M, and S states was compared between the two groups using Pearson’s Chi-Square test.

### Cell-Cell Communication Analysis Using CellChat

Cell-cell communication analysis was performed using the CellChat (RRID: SCR_021946) package in R. [43] NPC datasets were divided into two groups (UPS-high and UPS-low) to explore alterations in intercellular signaling, and each network was inferred separately. The ligand-receptor interactions were ranked based on interaction weight, and the patterns were compared between the two groups. Differential expression analysis was performed on ligand-receptor pairs with UPS-low cells designated as the reference group. Upregulated and downregulated signaling interactions were identified and mapped to ligand-receptor pairs. Bubble plots were used to visualize significantly upregulated and downregulated ligand-receptor interactions in UPS-high cancer cells. Network ranking plots were also created to compare the input and output signaling from the UPS-high and UPS-low cancer cells.

### Survival Analysis of the UPS Genes using GEPIA2

Survival analysis was performed using the GEPIA2 (RRID: SCR_026154) database (http://gepia2.cancer-pku.cn) [44] to evaluate the prognostic relevance of the 12 genes associated with UPS (CCT6A, CCT3, PSMA1, PSMC4, PSMA7, PSMA6, PSMA4, PSMD1, PSMD14, PSMD12, UCHL5, and CCT5). Each gene was individually assessed for its associations with overall survival across TCGA cancer types using the Survival Map module. Genes with log-rank test p < 0.05 were considered statistically significant. To assess the combined effect of the 12-gene set, Kaplan–Meier survival curves were generated using the Survival Analysis module for two datasets: the full TCGA pan-cancer data using all available tumor types and the TCGA head and neck squamous cell carcinoma (HNSC) data. Due to the paucity of available survival datasets for NPC, HNSC was used as the most specific available proxy. Patients were stratified into high- and low-expression groups based on median expression of the combined signature.

## Results

A meta-analysis of six publicly available datasets was performed to identify differentially expressed genes (DEGs) associated with nasopharyngeal carcinoma (NPC). This analysis integrated RNA-Seq and microarray datasets with a meta-regression model to correct batch effects. A total of 4004 genes were identified following multiple hypothesis correction (p < 0.01; Supplementary Materials 1). Genes with known interactions with the Epstein-Barr virus (EBV) were specifically selected to refine the findings further. Protein-protein interaction data for EBV-human interactions were retrieved from the VirHostNet database, yielding 548 interactions encompassing 14 EBV and 362 human proteins. Cross-referencing this dataset with the identified DEGs resulted in the retention of 85 genes. Figure 1 illustrates the PPI network of these genes and their corresponding EBV-associated interactions. Some of these EBV proteins interacting with the DEGs are involved in the EBV lytic cycle.

**Figure 1.**
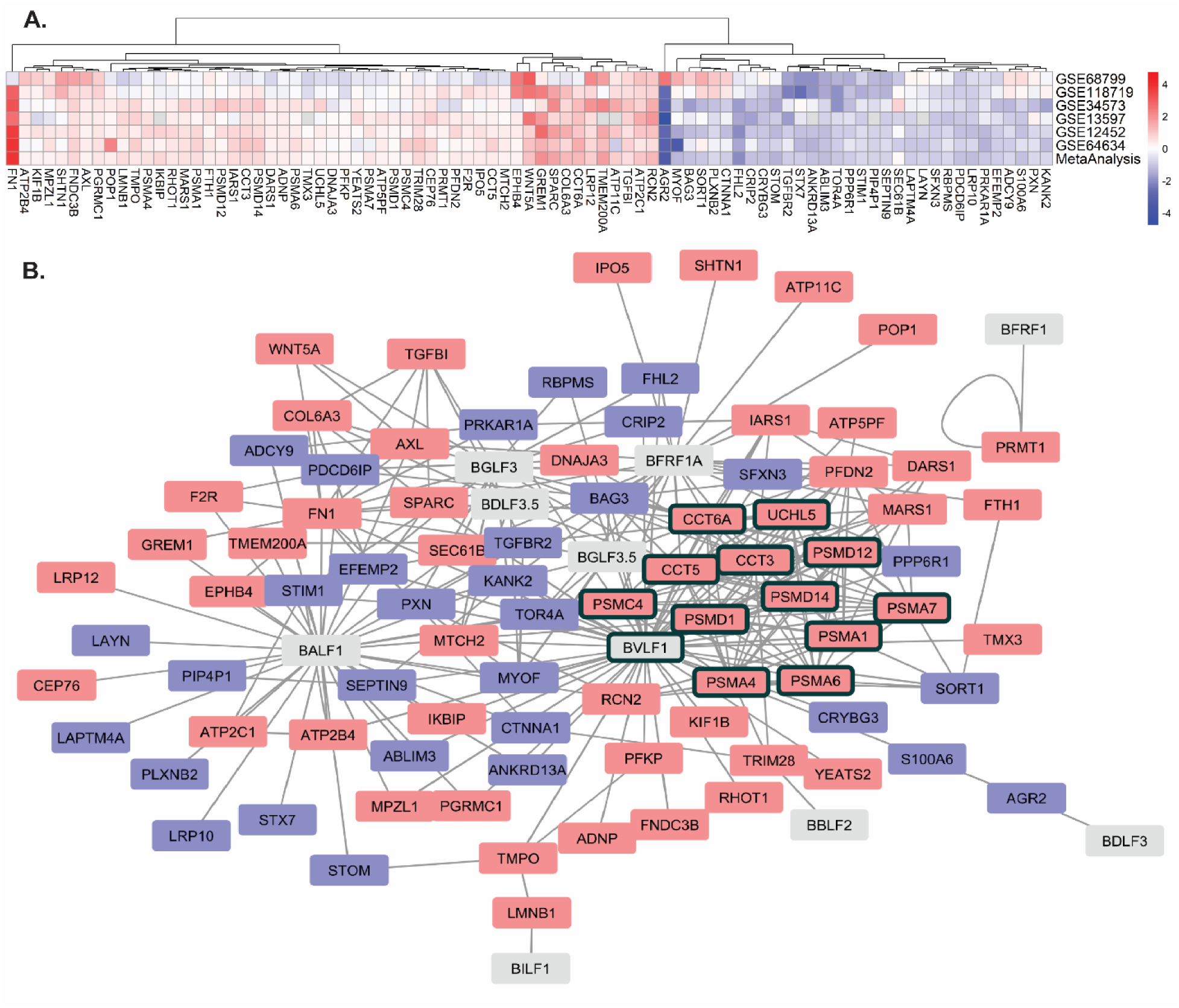
Differentially expressed genes (DEGs) and protein-protein interactions (PPI) network in nasopharyngeal carcinoma. **A.** A heatmap displaying the logFC changes of genes interacting with the Epstein-Barr virus. The X-axis represents the individual studies and the meta-analysis results while the Y-axis represents the genes. All genes are statistically significant after the meta-analysis at the level of p < 0.01 after Benjamini-Hochberg correction. **B.** A PPI network demonstrating the interaction between upregulated (red) and downregulated (blue) human proteins with EBV-related proteins (grey). Key functional clusters identified by the Molecular Complex Detection (MCODE) algorithm are outlined in black boxes, highlighting central hub genes relevant to NPC development.

Clusterization using MCODE on the PPI network identified several closely related proteins that may serve important functions in the pathogenesis of nasopharyngeal cancers. These genes are CCT6A, CCT3, PSMA1, PSMC4, PSMA7, PSMA6, PSMA4, PSMD1, PSMD14, PSMD12, UCHL5, and CCT5. Notably, all of these genes were influenced by several lytic genes, including BVLF1, BGLF3, BGLF3.5, BDLF3.5, and BFRF1A. Functional enrichment analysis of the clustered genes using the Reactome database and Gene Ontology (GO) Biological Process revealed that these genes are involved in the Antigen Presentation and the Ubiquitin-Proteasomal (UPS) Pathway (Table 2; Supplementary Materials 2). This finding establishes a link between late EBV gene expression and the regulation of this essential cellular degradation and protein turnover pathway, further highlighting its potential role in NPC tumorigenesis.

**Table 2.** Enrichment results of the MCODE clustered gene from the PPI network. The top two results from each database are shown.

| Databases | Term | Adjusted P-Value | Odds Ratio | Genes |
| --- | --- | --- | --- | --- |
| **Reactome Pathways** | Cross-presentation of Soluble Exogenous Antigens (Endosomes) | P < 0.001 | 1376.483 | PSMA6; PSMD12; PSMA4; PSMA1; PSMD14; PSMC4; PSMD1; PSMA7 |
|  | Regulation of Activated PAK-2p34 by Proteasome Mediated Degradation | P < 0.001 | 1376.483 | PSMA6; PSMD12; PSMA4; PSMA1; PSMD14; PSMC4; PSMD1; PSMA7 |
| **GO Biological Process** | Proteasomal Protein Catabolic Process (GO:0010498) | P < 0.001 | 186.57 | PSMA6; PSMD12; PSMA4; PSMA1; PSMD14; PSMC4; PSMD1; PSMA7 |
|  | Proteasome-Mediated Ubiquitin-Dependent Protein Catabolic Process (GO:0043161) | P < 0.001 | 125.54 | PSMA6; PSMD12; PSMA4; PSMA1; PSMD14; PSMC4; PSMD1; PSMA7 |

To further explore the role of the UPS-related genes in NPC tumorigenesis, we analyze several publicly available scRNA-Seq datasets from the GEO database. We use the HPV-negative oropharyngeal cancer dataset as a comparator due to its close anatomical proximity and similar normal histological characteristics to nasopharyngeal carcinoma. However, the HPV-negative cancer dataset is used solely as a reference for cancer cells in this region rather than implying histological equivalence between the malignancies. Initial copy number variations (CNV) analysis identified gross copy number abnormalities in the clusters initially identified as epithelial cells and sebocytes. These CNV alterations can be seen on chromosomes 1, 3, 4, 5, 6, 8, 11, 12, 14, and 17, reflecting the deranged cell division and proliferation typically found in cancer cells (Figure 2A). CNV alterations confirmed that these cells are indeed cancer cells, making them the focus of subsequent analysis. Comparing the EBV-related genes clustered by the MCODE algorithm showed statistically significantly higher expression of UPS-related genes in NPC cancer cells than in OPC (Figure 2C and 2D). This finding suggests NPC may utilize the UPS system more for cancer progression than OPC. Other cell types, including gamma delta T-cells, regulatory T-cells, and CD8+ T-cells, also showed high UPS activity, consistent with their established roles in antigen presentation and processing.

**Figure 2.**
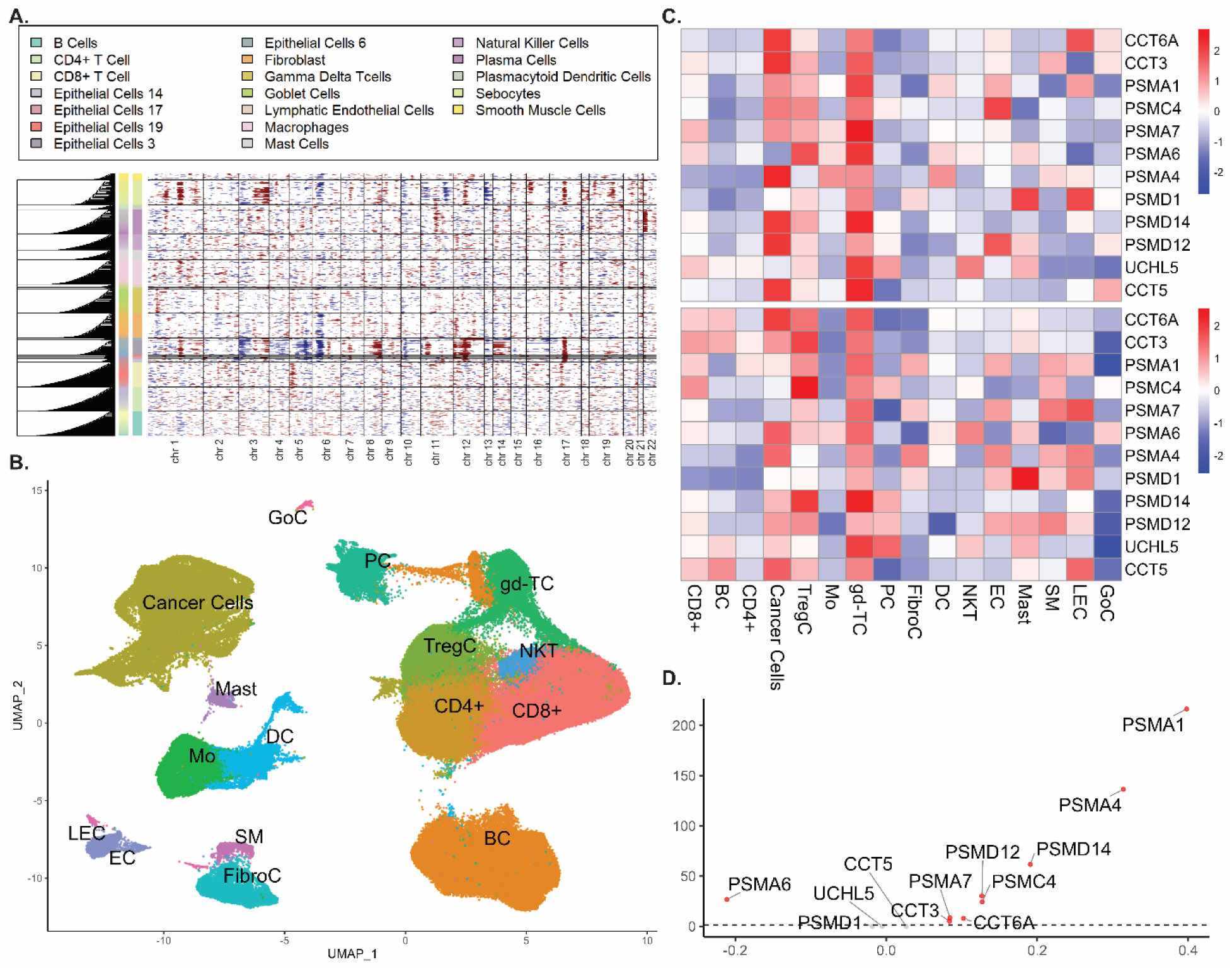
Genomic analysis and transcriptomic characterization of key genes between nasopharyngeal and oropharyngeal cancer cells. **A.** Copy number variation (CNV) analysis of cancer cells from the combined single-cell dataset. The heatmap displays CNV alterations across several specific cell types, notably the epithelial cell clusters (no 14, 17, 19, 3, and 6) and the sebocyte cell cluster. Cells with gross copy number alterations were subsequently classified as cancer cells. **B.** UMAP plot of single-cell clusters after relabeling. Each color represents a distinct cell type. **C.** Heatmap of the Ubiquitin-Proteasome System (UPS) related genes identified from the MCODE analysis. The heatmap compares gene expression between cell clusters in nasopharyngeal carcinoma (top) and oropharyngeal carcinoma (bottom), with red indicating upregulation and blue indicating downregulation. Note the high expression of the UPS-related genes in CD8+, Cancer Cells, Regulatory T-cells, and Gamma Delta T-cells, reflecting their increased UPS function. UPS-related genes are also more expressed in cancer cells from NPC than OPC. **D.** A volcano plot comparing the expression of the UPS-related genes between cancer cells from NPC and OPC. Significant genes are highlighted in red. The x-axis depicts the log-fold change (logFC), and the y-axis depicts the -log10 of the adjusted p-values. The dashed lines represent the line of no difference. **Cell Type Abbreviations:** **BC**: B-cells; **DC**: Dendritic cells; **CD4+:** CD4+ T-cells; **CD8+**: CD8+ T-cells; **EC**: Endothelial cells; **FibroC**: Fibroblasts; **gd-TC**: Gamma Delta T-cells; **LEC**: Lymphatic endothelial cells; **Mast**: Mast cells; **Mo**: Macrophages; **NKT**: Natural Killer T-cells; **PC**: Plasma cells; **SM**: Smooth muscle cells.

We performed a re-clustering analysis using UMAP on the NPC cancer cells (Figure 3A). We identified nine different clusters, implicating significant heterogeneity in their gene expression. To validate that these cells are indeed cancer cells, we overlay epithelial markers such as EPCAM, KRT8, KRT18, and CDH1 on these clusters (Figure 3B). The results supported that these heterogeneous clusters are indeed cancer cells. The heterogeneity of the cancer cells was further validated using enrichment of the marker genes for each cluster (Figure 3C). Our results showed that these cancer cells were enriched for different functions, with most of them enriched for immune-related signalling. Clusters 1, 3, 4, 6, 7, 8, and 10 were all enriched for different aspects of immune signaling reflecting the heavy interaction of NPC cancer cells with the resident immune system, which might explain the heavy immune infiltration usually found in NPC.

**Figure 3.**
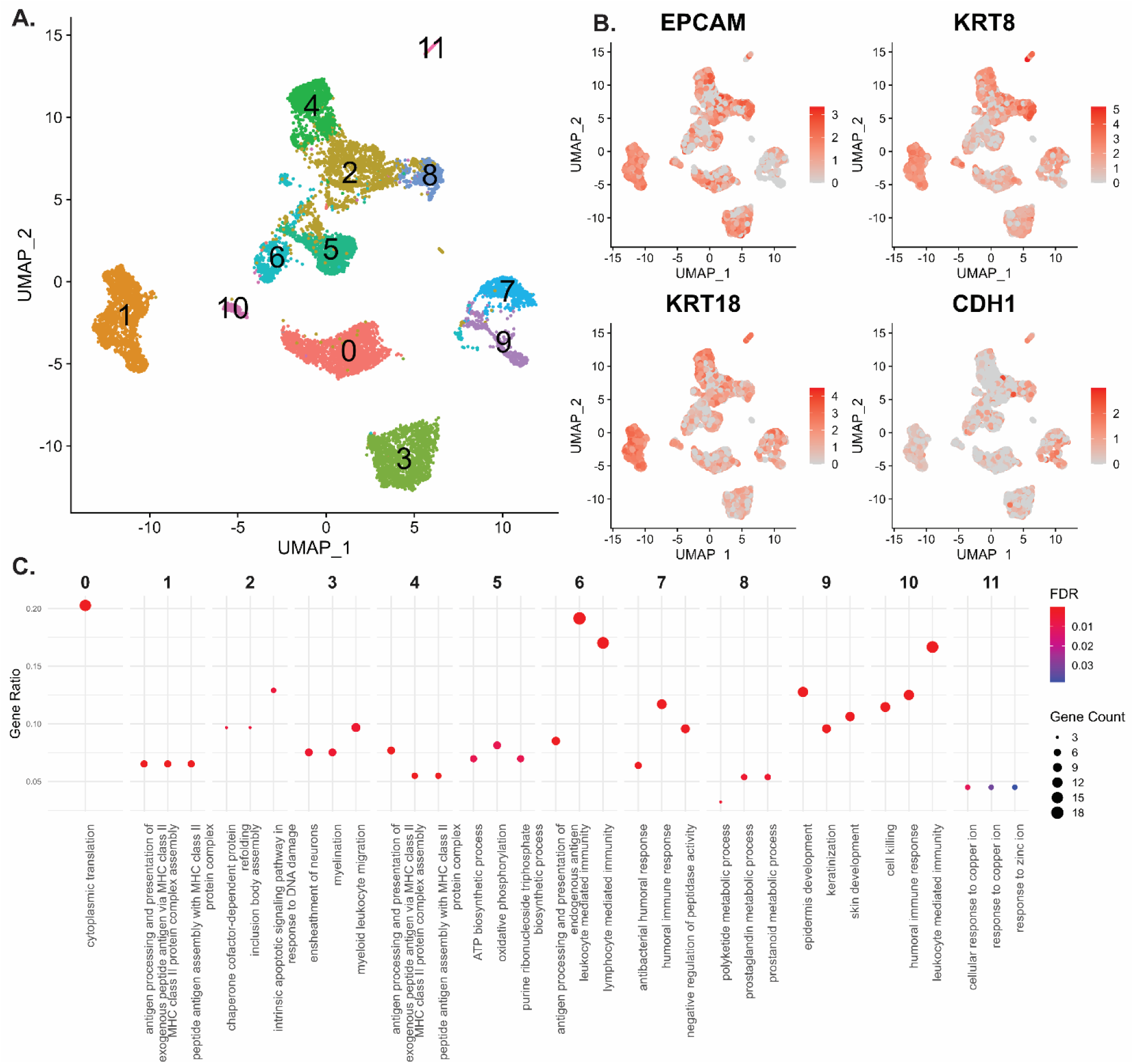
Re-clustering of cancer cells and enrichment of cancer cell clusters. **A.** UMAP plot displaying the reclustering of cancer cells from NPC. Distinct clusters can be seen, highlighting the heterogeneity of NPC cancer cells. **B.** Feature plot comparing the expression of Epithelial markers (EPCAM, KRT8, KRT18, and CDH1) across the different cancer cell clusters. Note that virtually all the cells in the tumor clusters express epithelial markers, validating their cancer identity. **C.** Dot plots comparing the enrichment of the top marker genes for each cluster. This plot identified that each cluster is associated with distinct characteristics. For example, clusters 6 and 10 were enriched for genes associated with the immune system, while those like cluster 9 are enriched for genes associated with skin development and keratinization.

We then further validated whether the upregulated UPS system previously identified has any roles in the immune signaling within NPC cancer cells. Using the AUCell package in R, we correlated the average expression of UPS-related gene expression with the pathway scores generated by AUCells (Figure 4A). We identified a significant positive correlation between the expression of UPS-related genes and immune-related pathways, such as T-cell-mediated immunity to tumor cells, and negative regulation of interferon and inflammatory response, to name a few. These findings suggested that UPS-High tumor cells are associated with less immune activation, leukocyte aggregation, and inflammatory responses. Taken together, these results suggest that UPS-High cells downregulate the inflammatory response and immune activity of the immune cells, helping them to survive in the dense immune environment.

**Figure 4.**
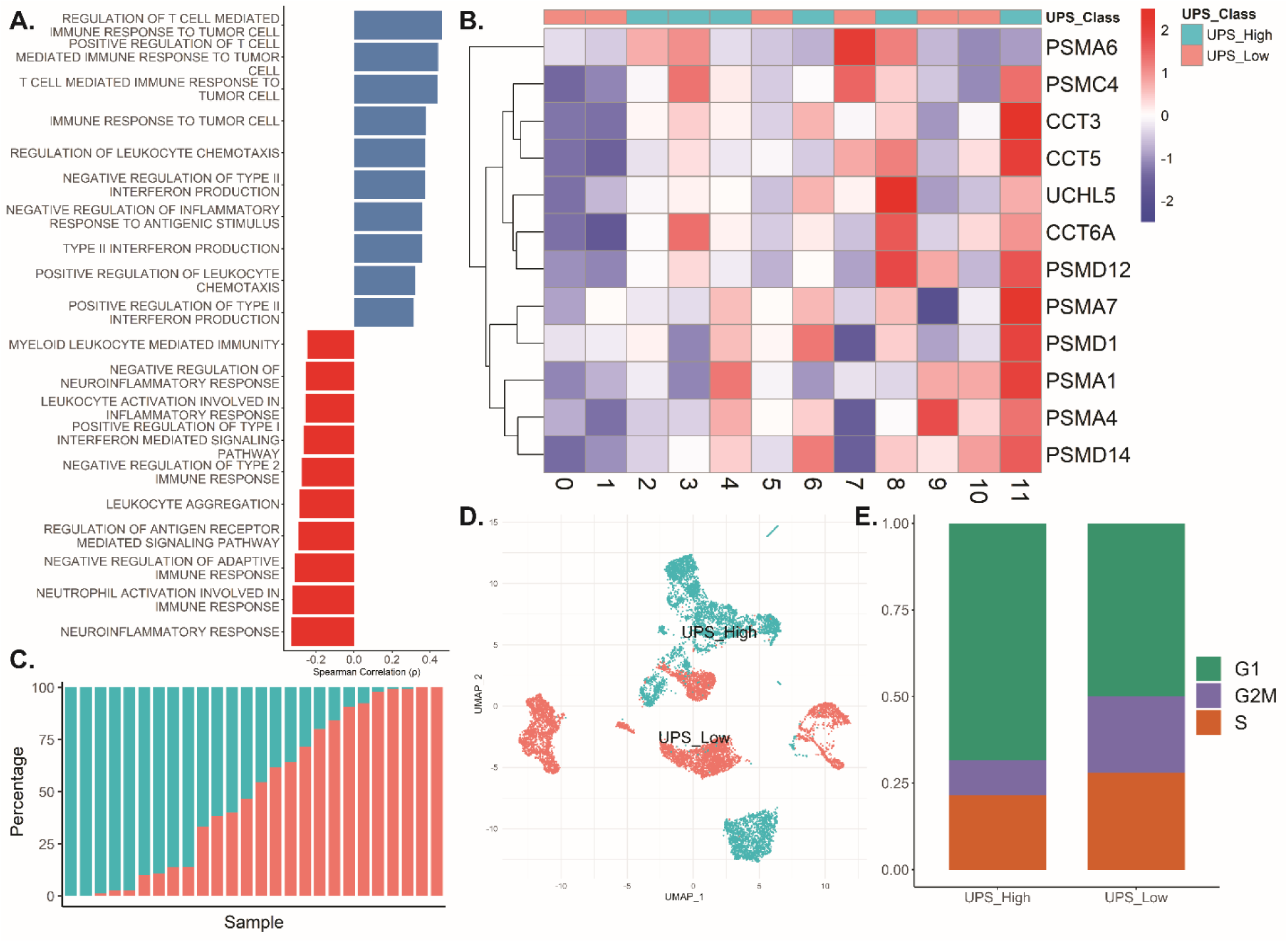
Characterization of UPS activity and its association with immune signaling, gene expression patterns, and cell cycle states. ***A.*** Bar plot showing Spearman correlation coefficients between the ubiquitin-proteasome system (UPS) expression signature and AUCell-derived activity scores of immune-related GO Biological Process terms across tumor cells. Note that a higher average expression of the UPS system is associated with a higher T-cell-mediated immune response and leukocyte chemotaxis. However, the higher UPS expression is also negatively correlated with the inflammatory response, neutrophil activation, and leukocyte activation. Perhaps suggesting that UPS-high tumor cells may be more visible to immune surveillance, but also downregulate inflammatory signalling to escape immune elimination. ***B.*** Heatmap comparing the expression of UPS-related genes across the different cancer cell clusters. Notably, clusters 2, 3, 4, 6, 8, and 11 showed increased UPS-related gene expression and are classified as UPS-High, while the remaining clusters are classified as UPS-Low. Red represents upregulated expression, while blue represents downregulated expression. ***C.*** Distribution of UPS-High and UPS-Low tumor cell proportions across patient samples. Each bar represents one patient, and the segments denote the relative percentage of UPS-High (blue) and UPS-Low (red) tumor cells. Note the inter-patient heterogeneity in UPS activity. ***D.*** UMAP plot showing the distribution of cancer cells after reclassification to UPS High and Low according to the expression of UPS-related genes. ***E.*** Bar plot comparing the proportion of cells in different cell cycle phases between UPS-High and UPS-Low groups. A markedly higher proportion of cells in the G1 phase is observed in the UPS-High group, with a corresponding lower proportion of cells in the S phase, indicating potential differences in cell cycle regulation. Proportions were tested using a chi-square test with p < 0.001.

A heatmap of UPS-related gene expression (Figure 4B) was created to compare their expression across cancer cell clusters. Note the significant heterogeneity of UPS-related gene expression on the cell clusters, potentially indicating the importance of UPS function in cancer cell progression. We then classified these clusters into UPS-High and UPS-Low using the median of their average expression as a cutoff. Clusters 2, 3, 4, 6, 8, and 11 were classified as UPS-High, while the rest were classified as UPS-Low.

We next assessed the distribution of UPS-High and UPS-Low tumor cells across individual patient samples to determine whether these subpopulations were restricted to specific patients (Figure 4C). Our analysis revealed that tumor cells with high or low UPS activity were distributed heterogeneously across the cohort. While some patients harbored predominantly UPS-High tumor cells, others exhibited mostly UPS-Low populations, with yet others showed a more balanced distribution between these two cell states. This observation supports the presence of substantial inter-patient heterogeneity in nasopharyngeal carcinoma, potentially driven by differential UPS activity, with implications for tumor growth dynamics, aggressiveness, and treatment potential.

We then examined the distribution of cancer cells in different cell cycle phases between UPS-High and UPS-Low groups (Figure 4E). The analysis revealed a higher proportion of cells in the S phase within the UPS-Low group, suggesting increased proliferative activity. Conversely, UPS-High cells exhibited a greater proportion in the G1 phase, indicative of a more quiescent state. These differences in proportion were tested using a chi-square test, yielding a statistically significant result (p < 0.001). These findings suggest that cancer cells in the UPS-High group may exhibit reduced proliferative signaling, possibly reflecting a shift toward a more quiescent, stem-like state.

To further investigate the role of UPS-related differences in cancer cell behavior, we analyzed the cancer cells from both groups and their interactions with immune cells within the tumor microenvironment. The analysis was restricted to immune cell interactions to assess differences in tumor-immune dynamics between UPS-High and UPS-Low cancer cells. Our analysis identified a marked difference in the number and strength of interactions between cancer cells and their microenvironment (Figure 5A). The cancer cells demonstrated mostly decreased self-interactions from the cancer cells to themselves and to other resident immune cells in the UPS-High group. However, the UPS-High group cells demonstrated increased communication strength compared to the UPS-Low group. Figure 5B illustrates the directional information flow of signaling to and from cancer cells, revealing enrichment of immune evasion–associated pathways (e.g., MIF, TNF, SIRP, and GALECTIN) in UPS-High cells. In contrast, UPS-Low cells exhibited significantly increased signaling in proliferation-related pathways such as WNT, FGF, and NOTCH.

**Figure 5.**
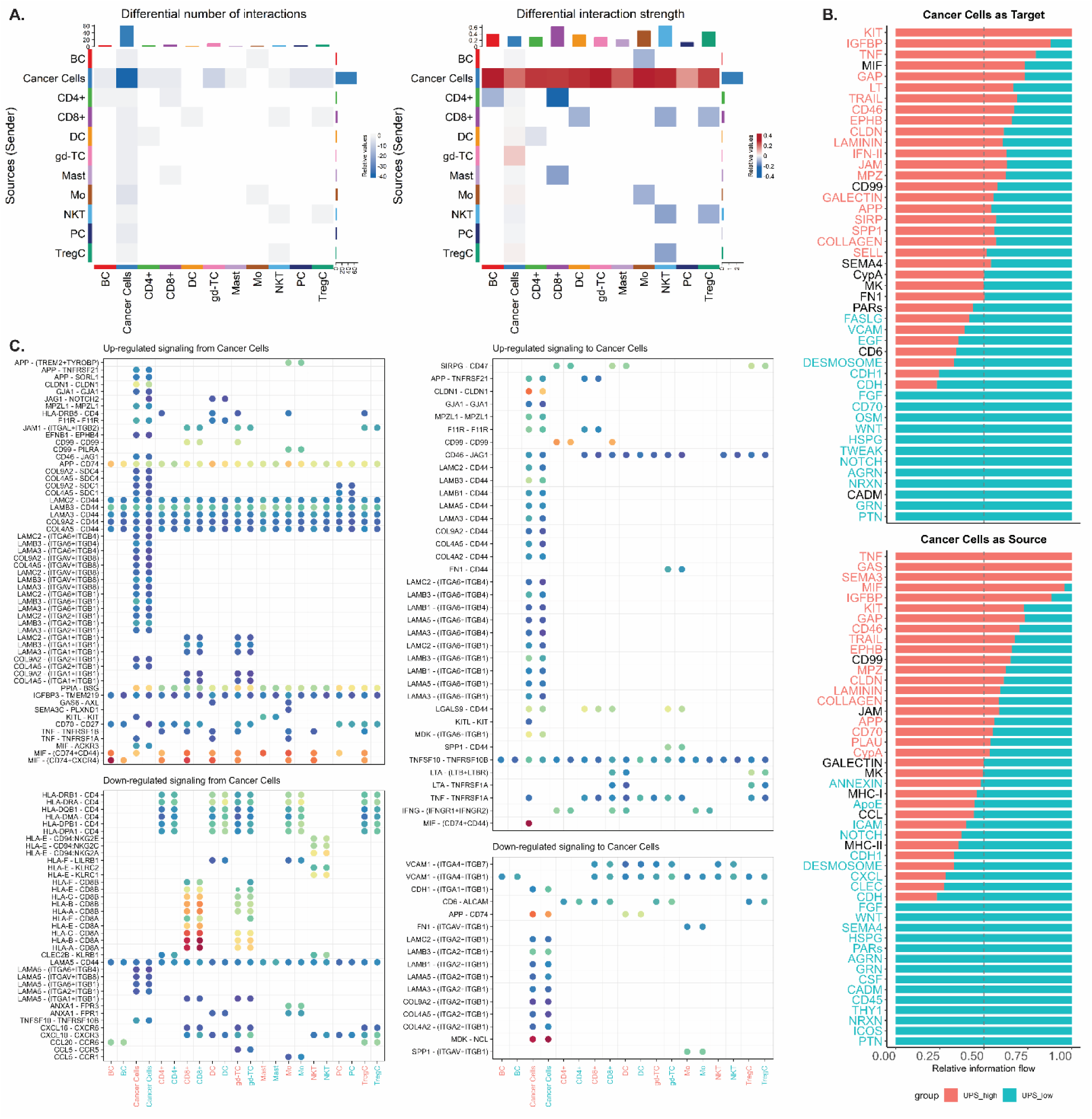
Cell interactions and differentially expressed ligand-receptor pairs between the UPS-High and UPS-Low cancer cells within the tumor microenvironment. ***A.*** Heatmap illustrating the number of interactions and interaction strength between cancer and resident immune cells. Red denotes higher interactions in the UPS-High group, while blue denotes higher interactions in the UPS-Low group. Note the marked decrease in the number of self-interactions between cancer cells and the higher interaction strength between cancer cells and the resident immune cells in the UPS-High tumor cells. ***B.*** Bar plots showing the relative information flow of signaling pathways where cancer cells act either as the target (top) or the source (bottom), stratified by UPS expression level. The relative information flow is defined as the ratio of signaling probability in one group divided by the sum of probabilities in both groups. Signaling pathways enriched in UPS-high cancer cells (red) include those related to immune evasion (MIF, TNF, GALECTIN) and cancer stemness (KIT). In contrast, UPS-low cells (blue) show higher activity in signaling pathways such as FGF, NOTCH, and WNT, implying association with cancer proliferation and growth. This shift reflects the potential for UPS-high tumor cells to adopt a more immune evasive and stem-like properties, while UPS-low cells may retain their high proliferation properties. ***C.*** Bubble plot of significantly upregulated and downregulated signaling from (left) and to (right) cancer cells. The size of each circle corresponds to the p-value, while the color intensity reflects the communication probability, with warmer colors indicating stronger interactions. The x-axis denotes the cell target of the interaction of the cancer cells from the UPS-High group (red) and the UPS-Low group (blue). Notable highlights include the significant upregulation of the MIF signaling pathway in the UPS-High cancer cells compared to the UPS-Low cancer cells. Notable highlights include the upregulation and downregulation of several specific Laminin signalling pathways in the UPS-High group, suggesting the variable effect Laminin signaling may have on NPC pathogenesis. However, note that the communication probabilities of Laminin signaling are quite low in both groups. **Cell Type Abbreviations:** **BC**: B-cells; **DC**: Dendritic cells; **CD4+:** CD4+ T-cells; **CD8+**: CD8+ T-cells; **EC**: Endothelial cells; **FibroC**: Fibroblast cells; **gd-TC**: Gamma Delta T-cells; **LEC**: Lymphatic endothelial cells; **Mast**: Mast cells; **Mo**: Macrophages; **NKT**: Natural Killer T-cells; **PC**: Plasma cells; **SM**: Smooth muscle cells.

Differential expression analysis of ligand-receptor pairs upregulated and downregulated from cancer cells (Figure 5C; Supplementary Materials 3) identified the downregulation of the MHC-I and MHC-II signaling pathways in the UPS-High NPC cancer cells, suggesting the increased immune evasion we previously identified during correlation analysis. Another notable pathway only identified in the UPS-High cells is MIF signaling, which is responsible for the negative regulation of immune activation. The MIF signaling is shown to be highly expressed in the UPS-High group with a high probability of signaling to all other resident immune cells, including CD4, CD8, Dendritic Cells, Macrophages, and Natural Killer T-Cells. A notable finding is the varying regulation of laminin signaling, with the ligands being selectively upregulated while some were downregulated. These suggest the complex function of Laminin signaling with the potential to support and inhibit cancer growth. Further details on the differential expression analysis results are shown in the supplementary materials.

Our previous analysis has shown that high UPS activity is associated with stemness features and immune evasion signaling. Therefore, we next aim to determine whether this phenotype translates into differences in patient survival. Due to the lack of NPC-specific survival datasets, we used the GEPIA2 platform to examine the prognostic value of the 12 UPS-related genes across the available cancer types from the TCGA datasets. Each gene was individually assessed in the Survival Map module, with several showing significant associations (p < 0.05) with overall survival (Figure 6A). We then analyzed the combined 12-gene signature in the full TCGA pan-cancer dataset (N = 9,502) and in the HNSC dataset (N = 518). We select this cancer as the closest available approximation to NPC. In both analyses, high UPS signature expression was associated with significantly poorer survival (HR = 1.5, p < 0.01 for pan-cancer; HR = 1.4, p = 0.02 for HNSC; Figure 6B). We acknowledge that these results will not fully answer their role in NPC, especially regarding EBV infection. However, our results support the hypothesis that elevated UPS activity is associated with worse survival and deserves further validation in NPC patients.

**Figure 6.**
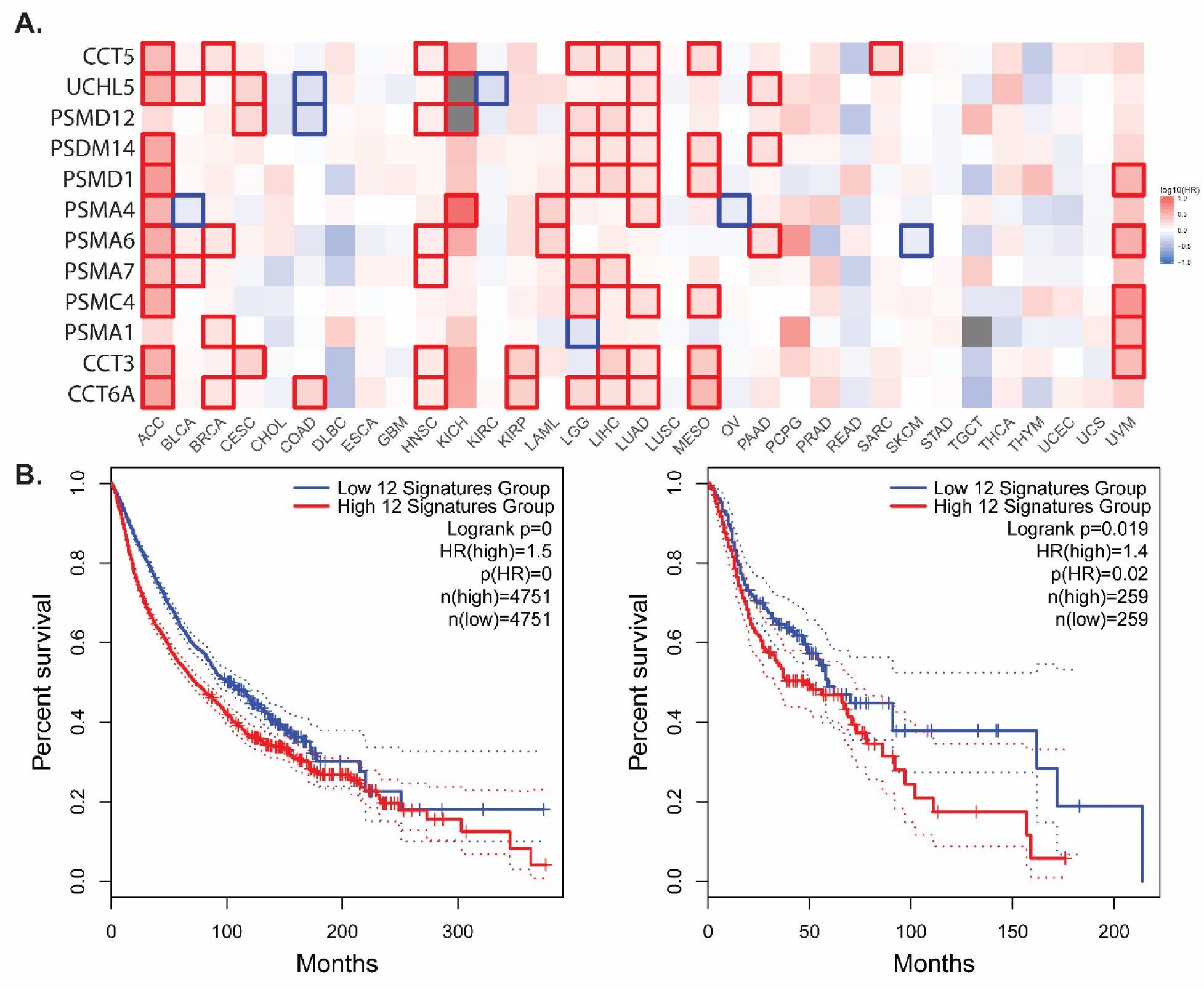
Prognostic relevance of the 12-gene UPS signature across cancers. ***A.*** Survival heatmap of each individual UPS gene (CCT6A, CCT3, PSMA1, PSMC4, PSMA7, PSMA6, PSMA4, PSMD1, PSMD14, PSMD12, UCHL5, and CCT5) across multiple cancer types from the TCGA pan-cancer dataset. The color scale represents the log10 hazard ratio for overall survival, with red indicating a higher risk associated with high gene expression and blue indicating a protective association. Significant associations (p < 0.05) are highlighted with red or blue boxes. ***B.*** Kaplan–Meier survival curves comparing patients with high versus low expression of the combined 12-gene UPS signature. **Left:** Pan-cancer dataset (N = 9,502; HR = 1.5, p < 0.01). **Right:** Head and neck squamous cell carcinoma dataset (N = 518; HR = 1.4, p = 0.02). Although NPC-specific data were unavailable, elevated UPS gene expression correlated with worse survival in pan-cancer and HNSC, warranting further validation.

## Discussion

In this study, we investigated and identified a complex interaction between EBV infection and the ubiquitin-proteasome system in NPC. Our findings provided further evidence on EBV’s major role in modulating several carcinogenesis signaling pathways. We identified that several genes associated with the EBV lytic cycle may upregulate UPS function with a subsequent decrease in cell proliferation, downregulation of genes associated with immune response, and upregulation of genes associated with cancer stemness and immune tolerance. By integrating publicly available transcriptomics datasets, we outline the effects of EBV on the development of NPC and explain its possible pathogenesis.

### DEGs expressed by NPCs are influenced primarily by EBV lytic proteins

Our analysis identified the enrichment of 85 EBV-interacting genes among the DEGs found through the meta-analysis. This finding highlights the importance of viral influence during the development of nasopharyngeal cancer. Previous studies have reported that EBV induces multiple alterations to the tumor microenvironment, especially due to its ability to infect multiple cells. Interestingly, contrary to previous findings that associated EBV carcinogenesis with the latent phase of the infection (Ahmed et al., 2022),our findings link the lytic cycle and the associated expression of the lytic proteins to EBV carcinogenesis. We found that of the DEGs identified to be EBV-related, almost all are influenced by EBV lytic proteins. Our findings align with several studies that found EBV exploits the host ubiquitination and proteasomal degradation processes to evade the immune system. [46,47] EBV also promoted the selective degradation of these proteins, targeting those proteins that promoted their cell growth and virus persistence. Therefore, the lytic phase of EBV infection might facilitate cancer persistence by facilitating immune escape and increasing the viral spread. It is important to note that several recent studies have highlighted the expression of EBV lytic proteins in NPC and other virus-associated tumours, implicating these EBV proteins in the oncogenic process and in immune evasion [5,48].

The UPS is essential for regulating protein turnover and antigen processing. The dysregulation of this system is implicated in various malignancies, including those associated with viral infections such as EBV. [49] Viruses use the UPS to degrade specific cellular targets to promote their replication. However, collateral damages usually occur, such as the degradation of p53 and the retinoblastoma tumor suppressor by the human papillomavirus (HPV) infection. [50] Using HPV-negative oropharyngeal carcinoma as a control of a similar location and histology, we identified that UPS-related genes are significantly upregulated. This reflects that although cancer cells have dysregulation of UPS activity, EBV infections cause further derangement to promote their persistence and cancer progression. The importance of UPS activation in cancers was also shown by studies exploring bortezomib, a proteasomal inhibitor, in enhancing cancer cell recognition by the immune system and reducing proliferation in multiple cancers, including NPC. [51–53]

### Cluster analysis revealed intratumoral heterogeneity of UPS functions

Given the heterogeneity of cancer cells, we reclustered the NPC cells to examine the expression of UPS-related genes further. Initially, we identified 12 distinct cancer cell clusters, each with different gene functions enriched. We identified that the majority of cancer cells were enriched with genes regulating the immune system. These supported previous findings of dense inflammatory infiltrate and interaction in nasopharyngeal cancers. [54–56] To evaluate the influence of UPS-related genes on the immune system, we correlate the average expression of UPS-related genes with the gene enrichment score generated using AUCells. We identified that high UPS expression is positively correlated with the regulation of T-cell-mediated response to tumor cells. These findings are supported by the main function of the UPS system in antigen presentation and its importance in cancer cell recognition. [57] However, we also identified that high UPS expression is negatively correlated with markers of immune response and inflammation. These were confirmed by previous studies that showed the UPS system as essential to regulating the immune response. [55,56] In nasopharyngeal cancers, we postulate that high UPS activity, although it may increase cancer cell recognition, also causes immune dysregulation, which may support their growth.

Using the average expression of UPS-related genes, we classified the cancer cells into two main clusters: UPS-High and UPS-Low. Previous studies have associated proteasomal dysfunctions with cancer growth. Consistent with previous findings, we identified statistically significant differences in the proportion of cells in the G1, G2/M, and S phases between the two groups. Higher proportions of cells in the S phase and lower proportions in the G1 phase were observed in the UPS-Low group, signifying higher proliferation. These findings suggest that cells with low UPS activity have a more rapid cell proliferation, presumably through the potential non-degradation of growth signals. Several studies have associated the deregulation of UPS activity with increased signaling activity of signaling pathways such as Wnt [58], NF-κB [59], PI3K/Akt [60], and Notch [61]. In our study, we also identified increased Wnt and FGF signaling to and from UPS-Low cancer cells (Figure 5B). The increased activity may lead to a pro-tumorigenic environment that supports cancer growth. Although cells in the UPS-High group exhibit lower proliferative signaling, they show increased activity in immune evasion pathways and expression of stemness-associated markers such as CD44. This phenotype suggests a subpopulation that may persist during treatment and contribute to eventual relapse. This highlights the relevance of proteasome inhibition in selectively targeting this immune-evasive, low proliferative, sub-population. However, the heterogeneous distribution of UPS-High and UPS-Low subsets across patients (Figure 4E) suggests that proteasome inhibition alone may not be sufficient. Effective treatment strategies may require combination therapies to eliminate both proliferative and immune-evasive tumor populations.

### Higher UPS activity is associated with immune evasion and tolerance

One of the hallmark features of cancer is the ability of cancer cells to evade immune detection. [62,63] The ability to evade the immune system is found in virus and non-virus-associated cancers. [64] However, carcinogenesis driven by viruses also adds another immune escape mechanism through viral immune evasion. [64,65] NPC was associated with marked downregulation of immune activity. [66,67] Our findings further linked these immune escape mechanisms with UPS activity, with marked downregulation of ligand-receptor pairs belonging to the MHC class I and II molecules in UPS-High cells. The downregulation of MHC class I and II signaling was observed between the cancer cells and the immune cells (CD4+, CD8+, γδ T-cells, macrophages, and regulatory T-cells), effectively making the cancer cells invisible to the immune system. Our findings are supported by previous studies identifying EBV proteins such as BDLF3 [66] and BILF1 [68], which promote the ubiquitination and degradation of MHC class I and II molecules. These observations suggest that EBV may substantially impair antigen presentation, likely through manipulation of the ubiquitin–proteasome system to enhance proteasomal degradation of immune recognition molecules.

We identified increased expression of macrophage inhibitory factor (MIF) signaling, and we identified that this signaling pathway is highly (and only) active in UPS-High cells. The ligand-receptor pairs MIF-CD74+CD44 and MIF-CD74+CXCR4 were found to be highly active in signaling between cancer cells with all the resident immune cells, including B Cells, CD4+ T-cells, CD8+ T-cells, dendritic cells, γδ T-cells, mast cells, macrophages, NK T-cells, plasma cells, and regulatory T-cells. MIF signaling has been implicated in various cancers and appears to be involved in almost all hallmarks of cancer activity. [69] A recent study by Chen et al. identified that MIF may be secreted using exosomes from NPC cells, promoting NPC metastasis. [70] MIF was also shown to have dual functions, with early activity eliciting an anti-tumor immune response. However, as the cancer bulk grows, it assumes a more pro-tumorigenic response, encouraging immune evasion and neovascularization. [71] Perhaps an interesting link between MIF and proteasomal activity can be found in research by Wang et al. on multiple myeloma (MM). [72] The inhibition of MIF resensitizes MM cells to the use of proteasomal inhibitors. Our findings suggest that increased UPS activity is also associated with increased MIF activity and reflects its potential in promoting immune tolerance to UPS-high nasopharyngeal cancer cells. These findings highlight a potential new combination of treatment targets for nasopharyngeal carcinoma.

Interestingly, we identified Galectin-9 (LGALS9) as an upregulated pathway in the nasopharyngeal cancer cells, with its interaction with CD44. Galectin-9 has emerged as a key immunoregulatory molecule, recognized in previous studies as a marker and mediator of T cell exhaustion. [73] Its interaction with CD44 contributes to immune tolerance by promoting the stability and suppressive function of regulatory T cells. [74] In a study by Kam et al. (Kam et al., 2025), elevated Galectin-9 expression in NPC tumor cells was shown to confer resistance to CD8 T cell-mediated cytotoxicity, highlighting its role in immune escape and poor immunotherapy response. Additionally, although not found in our data, the interaction of Galectin-9 and Tim3 has been the focus of recent research outlining its roles in immune escape and worse prognosis in nasopharyngeal cancers. [75,76] In our study, we identified that Galectin-9 signaling is upregulated in nasopharyngeal cancers in the UPS-high groups, supporting the role of this signaling in promoting immune escape.

### UPS activity is associated with both upregulation and downregulation of Laminin signaling

A particular intratumoral signaling pathways were downregulated in the UPS-High cell groups, the Laminin pathways. Laminins are large αβγ heterotrimeric glycoproteins that polymerise within the basement membrane to form a scaffold linking collagen to interact with cell surfaces. Laminins are crucial in cancers because they influence cancer growth, invasion, and metastasis. [77] We identified both upregulated and downregulated laminin ligand-receptor pairs (Figure 5C) in UPS-High NPC cells. The role of laminins in cancer was previously shown to be context-dependent [78], with both pro-tumorigenic effects during the loss and upregulation of laminin signaling. [79] For example, a study identified that the LAMC2 gene is overexpressed in esophageal cancers, and its abundance correlates with nodal metastasis and poor survival. [80] However, the same study also identified that inhibition of LAMA3 increased growth rates and migratory capacity of esophageal carcinomas, highlighting the dual roles of laminin. Multiple laminin-CD44 signaling was also observed, representing interactions between cancer cells and cancer cells with their microenvironment. CD44 is a well-established cancer stem cell marker implicated in tumor proliferation and treatment resistance through its interaction with the basement membrane and hyaluronic acid. [81–83] Furthermore, laminin signaling, particularly the expression of laminin subunits LAMB1 and laminin γ2, is recognized as a mediator of immune exclusion. A recent study in NPC identified that high LAMB1 expression is associated with poor progression-free survival and correlates with reduced infiltration of CD4 and CD8 T cells and reduced HLA expression, suggesting an immunosuppressive niche. [84] Although not specific to NPC, a study by Li et al. identified that tumor-derived laminin γ2 attenuated the response to anti–PD 1 therapies. [85] Thus, the concurrent upregulation of laminin–integrin signaling in UPS High cells likely contributes to a further immunosuppressive environment that reinforces resistance to therapy.

### Several classical growth factor signaling pathways are upregulated in UPS-Low cancer cells

The increased proportion of UPS-Low cancer cells in the replicative phase of the cell cycle (Figure 4D) may be caused by upregulated growth-promoting pathways. We identified two possible growth-promoting pathways, the FGF and the Wnt pathways. UPS-Low cancer cells were shown to upregulate FGF signalling, as shown by Figure 5B. FGF signaling derangements were shown to drive progression and resistance of cancer cells, such as lung, breast, and nasopharyngeal carcinomas. [86–89] A recent study exploring NPC identified that up to ∼70 % of tumours display high FGF1 transcripts, correlating with tumor growth through in vitro validations. [89] FGFR3 activation also influences macrophage phenotype, further complicating its role in NPC with its high immune infiltration. [90]. Another growth-promoting pathway identified is the canonical Wnt signalling. This pathway is well known in sustaining cancer-stem-cell pools and promotes cancer motility [91] and has also been identified to play a role in nasopharyngeal carcinoma pathogenesis [92,93]. Experimental over-expression of WNT10A stimulates migration, invasion, and self-renewal in squamous epithelium derived cancers and predicts poorer survival. [94] Together, these findings suggest that UPS-Low tumor cells exhibit enhanced growth-promoting signaling activity, particularly through the FGF and Wnt pathways, and likely represent the actively replicating compartment of the tumor, in contrast to the less proliferative, immune-evasive, and stem-like UPS-High population.

### High UPS activity is associated with worse prognosis in Pan-Cancer and HNSC datasets

Our survival analysis showed that elevated UPS gene expression is significantly associated with poorer outcomes in both the pan cancer and HNSC datasets. These findings were supported by other recent studies that showed the Protasome system is linked with reduced overall survival. A previous pan-cancer analysis has shown that expression of PSMA1 and PSMD11 was linked to unfavorable outcomes in more than 30% of cancer types tested. [95] Other authors have built upon these findings and created a prognostic model using UPS gene signatures in lung adenocarcinoma [96], thyroid carcinoma [97], and breast cancer. [98] These findings supported our results that the alterations of UPS, which are linked to stemness and immune evasion in NPC, are associated with adverse survival outcomes in many cancers. We acknowledge that without NPC-specific survival data, we cannot confirm these relationships. However, our findings should reinforce the UPS system as a potential biomarker and therapeutic target that deserves further experimental validation.

Our findings suggest a framework in which EBV drives NPC carcinogenesis (Figure 7). Although the traditional view emphasizes the latent EBV genes in NPC, our study identified a substantial influence of lytic phase proteins in tumorigenesis, with almost all DEGs regulated by the lytic proteins. We believe that while latent infections allow for long-term persistence of the virus, sporadic lytic activation is also essential for oncogenic transformation and immune evasion. One of the key cellular transformations caused by EBV is the manipulation of the UPS system. The dysfunctional UPS system promoted selective stabilization of oncogenic factors while promoting defective immune signaling and antigen presentation. Exploration of scRNA-seq data identified a marked downregulation in MHC-I and MHC-II signaling, coupled with the upregulation of other signaling pathways with the ability to promote immune escape, such as MIF. The UPS-high cells also upregulate and downregulate specific laminin signaling, highlighting their context-dependent functions. Several growth-promoting pathways, such as FGF and Wnt, were upregulated in the UPS-Low group, representing the highly proliferating tumor subpopulation. Taken together, the UPS system may contribute to differential regulation of NPC cancer cells and contribute to increased tumor persistence, proliferation, immune evasion, resistance to therapy, and adverse clinical outcomes.

**Figure 7.**
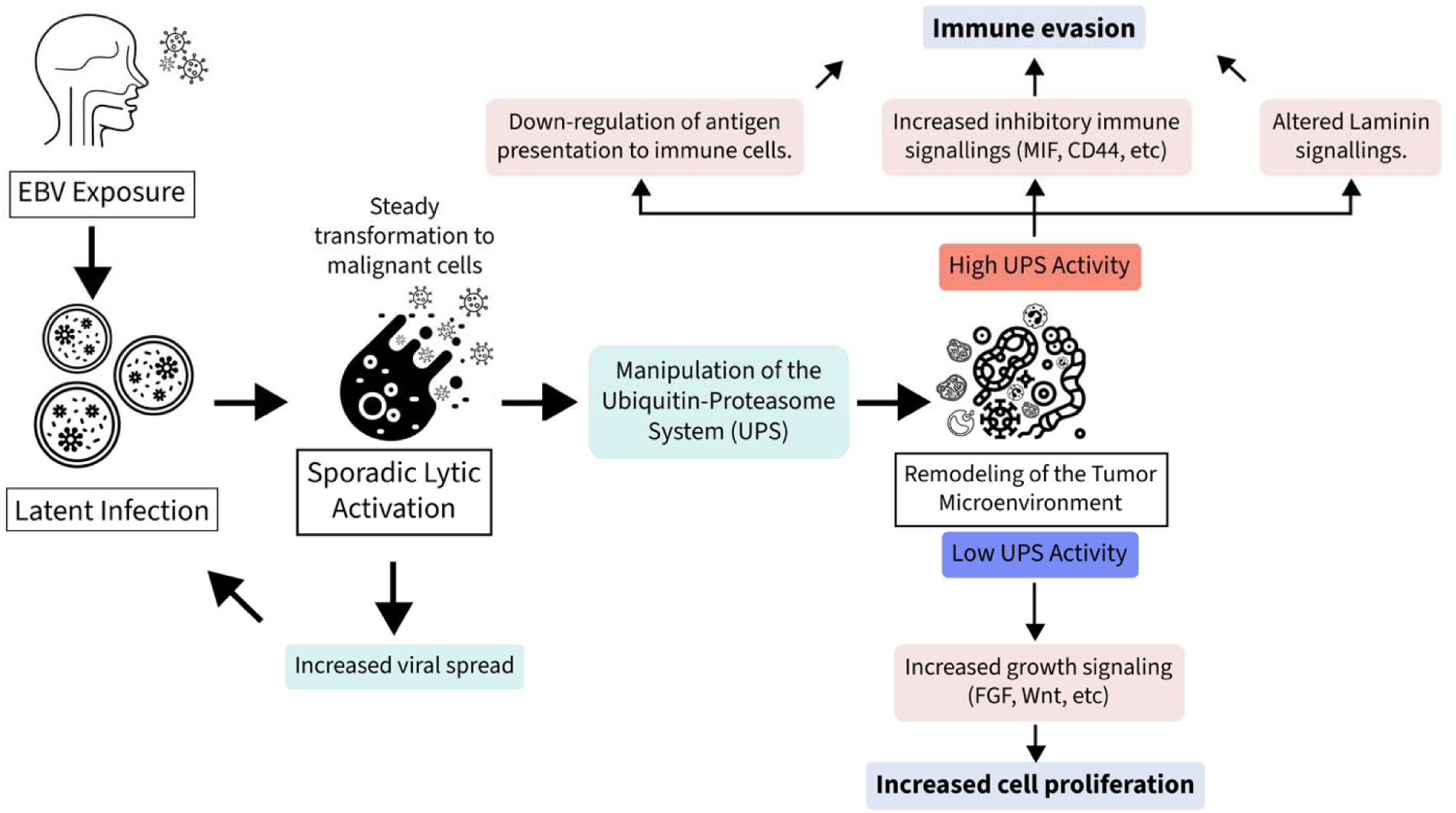
Proposed Framework of EBV-Driven NPC Progression. Upon EBV exposure, latent infection is established in epithelial cells, promoting malignant transformation. Sporadic lytic reactivation enhances viral spread and may contribute to additional oncogenic events. A central mechanism identified in our study is the manipulation of the ubiquitin– proteasome system (UPS), which drives remodeling of the tumor microenvironment. High UPS activity is associated with immune evasion, characterized by downregulation of antigen presentation and increased inhibitory signaling (e.g., MIF, CD44), along with altered laminin signaling. In contrast, low UPS activity is linked to increased tumor proliferation, supported by activation of growth-promoting pathways such as FGF and Wnt.

This study offers important insights into the role of the ubiquitin–proteasome system in EBV-associated nasopharyngeal carcinoma, but several limitations should be acknowledged. First, our conclusions are based solely on transcriptomic data, including bulk and single-cell RNA sequencing, without validation at the protein level. Therefore, our key findings, such as UPS activity, antigen presentation deficits, and ligand–receptor interactions (e.g., MIF, laminin, WNT, and FGF pathways), remain inferential. Second, cell–cell communication analysis using CellChat relies on probabilistic modeling of ligand–receptor co-expression and does not confirm actual physical or functional signaling, necessitating future validation through perturbation experiments. Third, the classification of UPS-High and UPS-Low tumor cells was based on relative (median) thresholds, which may not fully reflect real-world conditions. Fourth, while our survival analysis showed that high UPS gene expression is associated with poorer outcomes in both pan-cancer and HNSC datasets, the absence of NPC-specific survival data means these results should be interpreted cautiously. Lastly, although we observed substantial inter-patient heterogeneity, the number of available single-cell datasets remains limited, and the lack of integrated proteomics or clinical outcome data restricts generalizability. Future studies incorporating proteomic profiling, functional assays, and in vivo models will be essential to validate the therapeutic potential of targeting UPS activity and associated signaling phenotypes in NPC.

## Conclusion

This study highlights a framework of EBV-driven NPC progression, in which lytic-phase activation disrupts the ubiquitin–proteasome system (UPS), leading to altered immune signaling, reduced antigen presentation, increased stemness features, and context-dependent changes in pathways such as MIF, laminin, WNT, and FGF. These alterations appear to shape two distinct tumor cell phenotypes: UPS-High cells, which are less proliferative but more immune evasive, and UPS-Low cells, which are highly proliferative and enriched for growth-promoting signaling. Survival analysis further suggested that high UPS gene expression is associated with poorer outcomes in both pan-cancer and HNSC datasets, supporting its potential prognostic value; however, the absence of NPC-specific survival data means this relationship cannot be confirmed. While our findings integrate transcriptomic meta-analysis, single-cell resolution mapping, and exploratory survival analyses, they remain limited by the reliance on mRNA expression without direct proteomic validation, the lack of corroborative in vitro assays, and the probabilistic nature of inferred cell–cell communication results. Therefore, our conclusions should be interpreted cautiously and validated through future experiments, such as CRISPR-based gene editing or proteasomal inhibition assays, ideally in NPC-specific in vitro models, including organotypic co-cultures of tumor and immune cells. Nonetheless, the data support the hypothesis that targeting UPS activity and restoring antigen presentation may enhance immune recognition, while simultaneously disrupting pro-survival and stemness pathways, offering multiple therapeutic avenues for EBV-associated NPC.

## Declarations

### Ethics approval

Not applicable.

### Data and Code Availability

The public dataset used in the analysis are available from the GEO database from the following accession codes GSE68799, GSE118719 [24], GSE34573 [25], GSE13597 [26], GSE12452 [27], and GSE64634 [28] for bulk transcriptomics data and GSE150825, [34] GSE162025, [35] GSE150430, [36] GSE226620, [37] and GSE182227 [38] for the single-cell RNA Sequencing data. The code and script used in the analysis are available on figshare with doi:10.6084/m9.figshare.29900015 [99]

## Acknowledgments

The authors thanked Maranatha Christian University for providing the necessary facilities and funding during the creation of this research.

## Author Contributions

**Conceptualization**: Hana Ratnawati, Ardo Sanjaya;

**Methodology**: Ardo Sanjaya, Sascha Ott;

**Formal analysis**: Ardo Sanjaya, Sascha Ott;

**Data curation**: Ardo Sanjaya;

**Visualization**: Ardo Sanjaya, Aldrich Christiandy;

**Writing – original draft**: Hana Ratnawati, Ardo Sanjaya, Aldrich Christiandy;

**Writing – review and editing**: Hana Ratnawati, Ardo Sanjaya, Lawrence S. Young, Sascha Ott;

**Supervision**: Hana Ratnawati, Lawrence S. Young, Sascha Ott.

## References

1. Adham, M.; Kurniawan, A.N.; Muhtadi, A.I.; Roezin, A.; Hermani, B.; Gondhowiardjo, S.; Bing Tan, I.; Middeldorp, J.M. Nasopharyngeal Carcinoma in Indonesia: Epidemiology, Incidence, Signs, and Symptoms at Presentation. Chin J Cancer 2012, 31, 185–196, doi:10.5732/cjc.011.10328.

2. Chen, Y.P.; Chan, A.T.C.; Le, Q.T.; Blanchard, P.; Sun, Y.; Ma, J. Nasopharyngeal Carcinoma. The Lancet 2019, 394, 64–80.

3. Chang, E.T.; Ye, W.; Zeng, Y.X.; Adami, H.O. The Evolving Epidemiology of Nasopharyngeal Carcinoma. Cancer Epidemiology Biomarkers and Prevention 2021, 30, 1035–1047.

4. McKenzie, J.; El-Guindy, A. Epstein-Barr Virus Lytic Cycle Reactivation. In; 2015; pp. 237–261.

5. Yap, L.F.; Wong, A.K.C.; Paterson, I.C.; Young, L.S. Functional Implications of Epstein-Barr Virus Lytic Genes in Carcinogenesis. Cancers (Basel) 2022, 14, 5780, doi:10.3390/cancers14235780.

6. Dorothea, M.; Xie, J.; Yiu, S.P.T.; Chiang, A.K.S. Contribution of Epstein–Barr Virus Lytic Proteins to Cancer Hallmarks and Implications from Other Oncoviruses. Cancers (Basel) 2023, 15, 2120, doi:10.3390/cancers15072120.

7. Tsao, S.W.; Tsang, C.M.; Lo, K.W. Epstein-Barr Virus Infection and Nasopharyngeal Carcinoma. Philosophical Transactions of the Royal Society B: Biological Sciences 2017, 372, doi:10.1098/rstb.2016.0270.

8. Li, X.; Guo, Y.; Xiao, M.; Zhang, W. The Immune Escape Mechanism of Nasopharyngeal Carcinoma. The FASEB Journal 2023, 37, doi:10.1096/fj.202201628RR.

9. Yuan, L.; Zhong, L.; Krummenacher, C.; Zhao, Q.; Zhang, X. Epstein–Barr Virus-Mediated Immune Evasion in Tumor Promotion. Trends Immunol 2025, 46, 386–402, doi:10.1016/j.it.2025.03.007.

10. Chakravorty, S.; Afzali, B.; Kazemian, M. EBV-Associated Diseases: Current Therapeutics and Emerging Technologies. Front Immunol 2022, 13, doi:10.3389/fimmu.2022.1059133.

11. Zheng, X.; Huang, Y.; Li, K.; Luo, R.; Cai, M.; Yun, J. Immunosuppressive Tumor Microenvironment and Immunotherapy of Epstein–Barr Virus-Associated Malignancies. Viruses 2022, 14, 1017, doi:10.3390/v14051017.

12. Chen, J. Roles of the PI3K/Akt Pathway in Epstein-Barr Virus-Induced Cancers and Therapeutic Implications. World J Virol 2012, 1, 154, doi:10.5501/wjv.v1.i6.154.

13. Lo, A.K.-F.; Dawson, C.W.; Lung, H.L.; Wong, K.-L.; Young, L.S. The Role of EBV-Encoded LMP1 in the NPC Tumor Microenvironment: From Function to Therapy. Front Oncol 2021, 11, doi:10.3389/fonc.2021.640207.

14. Fang, W.; Zhang, J.; Hong, S.; Zhan, J.; Chen, N.; Qin, T.; Tang, Y.; Zhang, Y.; Kang, S.; Zhou, T.;, et al. EBV-Driven LMP1 and IFN-γ up-Regulate PD-L1 in Nasopharyngeal Carcinoma: Implications for Oncotargeted Therapy. Oncotarget 2014, 5, 12189–12202, doi:10.18632/oncotarget.2608.

15. Fukuda, M.; Longnecker, R. Epstein-Barr Virus Latent Membrane Protein 2A Mediates Transformation through Constitutive Activation of the Ras/PI3-K/Akt Pathway. J Virol 2007, 81, 9299–9306, doi:10.1128/JVI.00537-07.

16. Richardo, T.; Prattapong, P.; Ngernsombat, C.; Wisetyaningsih, N.; Iizasa, H.; Yoshiyama, H.; Janvilisri, T. Epstein-Barr Virus Mediated Signaling in Nasopharyngeal Carcinoma Carcinogenesis. Cancers (Basel) 2020, 12, 2441, doi:10.3390/cancers12092441.

17. Iwakiri, D. Epstein-Barr Virus-Encoded RNAs: Key Molecules in Viral Pathogenesis. Cancers (Basel) 2014, 6, 1615–1630, doi:10.3390/cancers6031615.

18. Samanta, M.; Iwakiri, D.; Takada, K. Epstein–Barr Virus-Encoded Small RNA Induces IL-10 through RIG-I-Mediated IRF-3 Signaling. Oncogene 2008, 27, 4150–4160, doi:10.1038/onc.2008.75.

19. Wang, M.; Yu, F.; Wu, W.; Wang, Y.; Ding, H.; Qian, L. Epstein-Barr Virus-Encoded MicroRNAs as Regulators in Host Immune Responses. Int J Biol Sci 2018, 14, 565–576, doi:10.7150/ijbs.24562.

20. Pei, Y.; Robertson, E.S. The Central Role of the Ubiquitin–Proteasome System in EBV-Mediated Oncogenesis. Cancers (Basel) 2022, 14, 611, doi:10.3390/cancers14030611.

21. Wang, L.; Ning, S. New Look of EBV LMP1 Signaling Landscape. Cancers (Basel) 2021, 13, 5451, doi:10.3390/cancers13215451.

22. Oswald, J.; Constantine, M.; Adegbuyi, A.; Omorogbe, E.; Dellomo, A.J.; Ehrlich, E.S. E3 Ubiquitin Ligases in Gammaherpesviruses and HIV: A Review of Virus Adaptation and Exploitation. Viruses 2023, 15, 1935, doi:10.3390/v15091935.

23. Lui, W.-Y.; Bharti, A.; Wong, N.-H.M.; Jangra, S.; Botelho, M.G.; Yuen, K.-S.; Jin, D.-Y. Suppression of CGAS- and RIG-I-Mediated Innate Immune Signaling by Epstein-Barr Virus Deubiquitinase BPLF1. PLoS Pathog 2023, 19, e1011186, doi:10.1371/journal.ppat.1011186.

24. Lin, C.; Zong, J.; Lin, W.; Wang, M.; Xu, Y.; Zhou, R.; Lin, S.; Guo, Q.; Chen, H.; Ye, Y.;, et al. EBV-MiR-BART8-3p Induces Epithelial-Mesenchymal Transition and Promotes Metastasis of Nasopharyngeal Carcinoma Cells through Activating NF-ΚB and Erk1/2 Pathways. Journal of Experimental & Clinical Cancer Research 2018, 37, 283, doi:10.1186/s13046-018-0953-6.

25. Hu, C.; Wei, W.; Chen, X.; Woodman, C.B.; Yao, Y.; Nicholls, J.M.; Joab, I.; Sihota, S.K.; Shao, J.-Y.; Derkaoui, K.D.;, et al. A Global View of the Oncogenic Landscape in Nasopharyngeal Carcinoma: An Integrated Analysis at the Genetic and Expression Levels. PLoS One 2012, 7, e41055, doi:10.1371/journal.pone.0041055.

26. Bose, S.; Yap, L.; Fung, M.; Starzcynski, J.; Saleh, A.; Morgan, S.; Dawson, C.; Chukwuma, M.B.; Maina, E.; Buettner, M.;, et al. The ATM Tumour Suppressor Gene Is Down-regulated in EBV-associated Nasopharyngeal Carcinoma. J Pathol 2009, 217, 345–352, doi:10.1002/path.2487.

27. Dodd, L.E.; Sengupta, S.; Chen, I.-H.; den Boon, J.A.; Cheng, Y.-J.; Westra, W.; Newton, M.A.; Mittl, B.F.; McShane, L.; Chen, C.-J.;, et al. Genes Involved in DNA Repair and Nitrosamine Metabolism and Those Located on Chromosome 14q32 Are Dysregulated in Nasopharyngeal Carcinoma. *Cancer Epidemiology*, Biomarkers & Prevention 2006, 15, 2216–2225, doi:10.1158/1055-9965.EPI-06-0455.

28. Bo, H.; Gong, Z.; Zhang, W.; Li, X.; Zeng, Y.; Liao, Q.; Chen, P.; Shi, L.; Lian, Y.; Jing, Y.;, et al. Upregulated Long Non-Coding RNA AFAP1-AS1 Expression Is Associated with Progression and Poor Prognosis of Nasopharyngeal Carcinoma. Oncotarget 2015, 6, 20404–20418, doi:10.18632/oncotarget.4057.

29. Ritchie, M.E.; Phipson, B.; Wu, D.; Hu, Y.; Law, C.W.; Shi, W.; Smyth, G.K. Limma Powers Differential Expression Analyses for RNA-Sequencing and Microarray Studies. Nucleic Acids Res 2015, 43, e47, doi:10.1093/nar/gkv007.

30. Viechtbauer, W. Conducting Meta-Analyses in *R* with the Metafor Package. J Stat Softw 2010, 36, doi:10.18637/jss.v036.i03.

31. Shannon, P.; Markiel, A.; Ozier, O.; Baliga, N.S.; Wang, J.T.; Ramage, D.; Amin, N.; Schwikowski, B.; Ideker, T. Cytoscape: A Software Environment for Integrated Models of Biomolecular Interaction Networks. Genome Res 2003, 13, 2498–2504, doi:10.1101/gr.1239303.

32. Bader, G.D.; Hogue, C.W. An Automated Method for Finding Molecular Complexes in Large Protein Interaction Networks. BMC Bioinformatics 2003, 4, 2, doi:10.1186/1471-2105-4-2.

33. Milacic, M.; Beavers, D.; Conley, P.; Gong, C.; Gillespie, M.; Griss, J.; Haw, R.; Jassal, B.; Matthews, L.; May, B.;, et al. The Reactome Pathway Knowledgebase 2024. Nucleic Acids Res 2024, 52, D672–D678, doi:10.1093/nar/gkad1025.

34. Gong, L.; Kwong, D.L.-W.; Dai, W.; Wu, P.; Li, S.; Yan, Q.; Zhang, Y.; Zhang, B.; Fang, X.; Liu, L.;, et al. Comprehensive Single-Cell Sequencing Reveals the Stromal Dynamics and Tumor-Specific Characteristics in the Microenvironment of Nasopharyngeal Carcinoma. Nat Commun 2021, 12, 1540, doi:10.1038/s41467-021-21795-z.

35. Liu, Y.; He, S.; Wang, X.-L.; Peng, W.; Chen, Q.-Y.; Chi, D.-M.; Chen, J.-R.; Han, B.-W.; Lin, G.-W.; Li, Y.-Q.;, et al. Tumour Heterogeneity and Intercellular Networks of Nasopharyngeal Carcinoma at Single Cell Resolution. Nat Commun 2021, 12, 741, doi:10.1038/s41467-021-21043-4.

36. Chen, Y.-P.; Yin, J.-H.; Li, W.-F.; Li, H.-J.; Chen, D.-P.; Zhang, C.-J.; Lv, J.-W.; Wang, Y.- Q.; Li, X.-M.; Li, J.-Y.;, et al. Single-Cell Transcriptomics Reveals Regulators Underlying Immune Cell Diversity and Immune Subtypes Associated with Prognosis in Nasopharyngeal Carcinoma. Cell Res 2020, 30, 1024–1042, doi:10.1038/s41422-020-0374-x.

37. Cha, J.; Kim, D.H.; Kim, G.; Cho, J.-W.; Sung, E.; Baek, S.; Hong, M.H.; Kim, C.G.; Sim, N.S.; Hong, H.J.;, et al. Single-Cell Analysis Reveals Cellular and Molecular Factors Counteracting HPV-Positive Oropharyngeal Cancer Immunotherapy Outcomes. J Immunother Cancer 2024, 12, doi:10.1136/jitc-2023-008667.

38. Puram, S. V; Mints, M.; Pal, A.; Qi, Z.; Reeb, A.; Gelev, K.; Barrett, T.F.; Gerndt, S.; Liu, P.; Parikh, A.S.;, et al. Cellular States Are Coupled to Genomic and Viral Heterogeneity in HPV-Related Oropharyngeal Carcinoma. Nat Genet 2023, 55, 640–650, doi:10.1038/s41588-023-01357-3.

39. Hao, Y.; Stuart, T.; Kowalski, M.H.; Choudhary, S.; Hoffman, P.; Hartman, A.; Srivastava, A.; Molla, G.; Madad, S.; Fernandez-Granda, C.;, et al. Dictionary Learning for Integrative, Multimodal and Scalable Single-Cell Analysis. Nat Biotechnol 2024, 42, 293–304, doi:10.1038/s41587-023-01767-y.

40. Korsunsky, I.; Millard, N.; Fan, J.; Slowikowski, K.; Zhang, F.; Wei, K.; Baglaenko, Y.; Brenner, M.; Loh, P.; Raychaudhuri, S. Fast, Sensitive and Accurate Integration of Single-Cell Data with Harmony. Nat Methods 2019, 16, 1289–1296, doi:10.1038/s41592-019-0619-0.

41. Kuleshov, M. V.; Jones, M.R.; Rouillard, A.D.; Fernandez, N.F.; Duan, Q.; Wang, Z.; Koplev, S.; Jenkins, S.L.; Jagodnik, K.M.; Lachmann, A.;, et al. Enrichr: A Comprehensive Gene Set Enrichment Analysis Web Server 2016 Update. Nucleic Acids Res 2016, 44, W90–W97, doi:10.1093/nar/gkw377.

42. Tickle, T.; Tirosh, I.; Georgescu, C.; Brown, M.; Haas, B. InferCNV of the Trinity CTAT Project. 2019.

43. Jin, S.; Plikus, M. V.; Nie, Q. CellChat for Systematic Analysis of Cell–Cell Communication from Single-Cell Transcriptomics. Nat Protoc 2025, 20, 180–219, doi:10.1038/s41596-024-01045-4.

44. Tang, Z.; Kang, B.; Li, C.; Chen, T.; Zhang, Z. GEPIA2: An Enhanced Web Server for Large-Scale Expression Profiling and Interactive Analysis. Nucleic Acids Res 2019, 47, W556–W560, doi:10.1093/nar/gkz430.

45. Ahmed, N.; Abusalah, M.A.H.A.; Farzand, A.; Absar, M.; Yusof, N.Y.; Rabaan, A.A.; AlSaihati, H.; Alshengeti, A.; Alwarthan, S.; Alsuwailem, H.S.;, et al. Updates on Epstein– Barr Virus (EBV)-Associated Nasopharyngeal Carcinoma: Emphasis on the Latent Gene Products of EBV. Medicina (B Aires) 2022, 59, 2, doi:10.3390/medicina59010002.

46. Silva, J. de M.; Alves, C.E. de C.; Pontes, G.S. Epstein-Barr Virus: The Mastermind of Immune Chaos. Front Immunol 2024, 15, doi:10.3389/fimmu.2024.1297994.

47. Quinn, L.L.; Williams, L.R.; White, C.; Forrest, C.; Zuo, J.; Rowe, M. The Missing Link in Epstein-Barr Virus Immune Evasion: The BDLF3 Gene Induces Ubiquitination and Downregulation of Major Histocompatibility Complex Class I (MHC-I) and MHC-II. J Virol 2016, 90, 356–367, doi:10.1128/JVI.02183-15.

48. Deng, Y.; Münz, C. Roles of Lytic Viral Replication and Co-Infections in the Oncogenesis and Immune Control of the Epstein–Barr Virus. Cancers (Basel) 2021, 13, 2275, doi:10.3390/cancers13092275.

49. Zhou, X.; Xu, R.; Wu, Y.; Zhou, L.; Xiang, T. The Role of Proteasomes in Tumorigenesis. Genes Dis 2024, 11, 101070, doi:10.1016/j.gendis.2023.06.037.

50. Moody, C.A.; Laimins, L.A. Human Papillomavirus Oncoproteins: Pathways to Transformation. Nat Rev Cancer 2010, 10, 550–560, doi:10.1038/nrc2886.

51. Jiang, G.-M.; Wang, H.-S.; Du, J.; Ma, W.-F.; Wang, H.; Qiu, Y.; Zhang, Q.-G.; Xu, W.; Liu, H.-F.; Liang, J.-P. Bortezomib Relieves Immune Tolerance in Nasopharyngeal Carcinoma via STAT1 Suppression and Indoleamine 2,3-Dioxygenase Downregulation. Cancer Immunol Res 2017, 5, 42–51, doi:10.1158/2326-6066.CIR-16-0102.

52. Sogbein, O.; Paul, P.; Umar, M.; Chaari, A.; Batuman, V.; Upadhyay, R. Bortezomib in Cancer Therapy: Mechanisms, Side Effects, and Future Proteasome Inhibitors. Life Sci 2024, 358, 123125, doi:10.1016/j.lfs.2024.123125.

53. Yarza, R.; Bover, M.; Agulló-Ortuño, M.T.; Iglesias-Docampo, L.C. Current Approach and Novel Perspectives in Nasopharyngeal Carcinoma: The Role of Targeting Proteasome Dysregulation as a Molecular Landmark in Nasopharyngeal Cancer. Journal of Experimental & Clinical Cancer Research 2021, 40, 202, doi:10.1186/s13046-021-02010-9.

54. Forder, A.; Stewart, G.L.; Telkar, N.; Lam, W.L.; Garnis, C. New Insights into the Tumour Immune Microenvironment of Nasopharyngeal Carcinoma. Current Research in Immunology 2022, 3, 222–227, doi:10.1016/j.crimmu.2022.08.009.

55. Beck, D.B.; Werner, A.; Kastner, D.L.; Aksentijevich, I. Disorders of Ubiquitylation: Unchained Inflammation. Nat Rev Rheumatol 2022, 18, 435–447, doi:10.1038/s41584-022-00778-4.

56. Goetzke, C.C.; Ebstein, F.; Kallinich, T. Role of Proteasomes in Inflammation. J Clin Med 2021, 10, 1783, doi:10.3390/jcm10081783.

57. Çetin, G.; Klafack, S.; Studencka-Turski, M.; Krüger, E.; Ebstein, F. The Ubiquitin– Proteasome System in Immune Cells. Biomolecules 2021, 11, 60, doi:10.3390/biom11010060.

58. Park, H.-B.; Kim, J.-W.; Baek, K.-H. Regulation of Wnt Signaling through Ubiquitination and Deubiquitination in Cancers. Int J Mol Sci 2020, 21, 3904, doi:10.3390/ijms21113904.

59. Zhou, X.; Xu, R.; Wu, Y.; Zhou, L.; Xiang, T. The Role of Proteasomes in Tumorigenesis. Genes Dis 2024, 11, 101070, doi:10.1016/j.gendis.2023.06.037.

60. Jin, J.; He, J.; Li, X.; NI, X.; Jin, X. The Role of Ubiquitination and Deubiquitination in PI3K/AKT/MTOR Pathway: A Potential Target for Cancer Therapy. Gene 2023, 889, 147807, doi:10.1016/j.gene.2023.147807.

61. Shi, Q.; Xue, C.; Zeng, Y.; Yuan, X.; Chu, Q.; Jiang, S.; Wang, J.; Zhang, Y.; Zhu, D.; Li, L. Notch Signaling Pathway in Cancer: From Mechanistic Insights to Targeted Therapies. Signal Transduct Target Ther 2024, 9, 128, doi:10.1038/s41392-024-01828-x.

62. Hanahan, D. Hallmarks of Cancer: New Dimensions. Cancer Discov 2022, 12, 31–46, doi:10.1158/2159-8290.CD-21-1059.

63. Galassi, C.; Chan, T.A.; Vitale, I.; Galluzzi, L. The Hallmarks of Cancer Immune Evasion. Cancer Cell 2024, 42, 1825–1863, doi:10.1016/j.ccell.2024.09.010.

64. Roetman, J.J.; Apostolova, M.K.I.; Philip, M. Viral and Cellular Oncogenes Promote Immune Evasion. Oncogene 2022, 41, 921–929, doi:10.1038/s41388-021-02145-1.

65. Cao, S.; Wylie, K.M.; Wyczalkowski, M.A.; Karpova, A.; Ley, J.; Sun, S.; Mashl, R.J.; Liang, W.-W.; Wang, X.; Johnson, K.;, et al. Dynamic Host Immune Response in Virus-Associated Cancers. Commun Biol 2019, 2, 109, doi:10.1038/s42003-019-0352-3.

66. Quinn, L.L.; Williams, L.R.; White, C.; Forrest, C.; Zuo, J.; Rowe, M. The Missing Link in Epstein-Barr Virus Immune Evasion: The BDLF3 Gene Induces Ubiquitination and Downregulation of Major Histocompatibility Complex Class I (MHC-I) and MHC-II. J Virol 2016, 90, 356–367, doi:10.1128/JVI.02183-15.

67. Zuo, J.; Currin, A.; Griffin, B.D.; Shannon-Lowe, C.; Thomas, W.A.; Ressing, M.E.; Wiertz, E.J.H.J.; Rowe, M. The Epstein-Barr Virus G-Protein-Coupled Receptor Contributes to Immune Evasion by Targeting MHC Class I Molecules for Degradation. PLoS Pathog 2009, 5, e1000255, doi:10.1371/journal.ppat.1000255.

68. Zuo, J.; Quinn, L.L.; Tamblyn, J.; Thomas, W.A.; Feederle, R.; Delecluse, H.-J.; Hislop, A.D.; Rowe, M. The Epstein-Barr Virus-Encoded BILF1 Protein Modulates Immune Recognition of Endogenously Processed Antigen by Targeting Major Histocompatibility Complex Class I Molecules Trafficking on Both the Exocytic and Endocytic Pathways. J Virol 2011, 85, 1604–1614, doi:10.1128/JVI.01608-10.

69. Mora Barthelmess, R.; Stijlemans, B.; Van Ginderachter, J.A. Hallmarks of Cancer Affected by the MIF Cytokine Family. Cancers (Basel) 2023, 15, 395, doi:10.3390/cancers15020395.

70. Chen, W.; Zuo, F.; Zhang, K.; Xia, T.; Lei, W.; Zhang, Z.; Bao, L.; You, Y. Exosomal MIF Derived From Nasopharyngeal Carcinoma Promotes Metastasis by Repressing Ferroptosis of Macrophages. Front Cell Dev Biol 2021, 9, doi:10.3389/fcell.2021.791187.

71. Noe, J.T.; Mitchell, R.A. MIF-Dependent Control of Tumor Immunity. Front Immunol 2020, 11, doi:10.3389/fimmu.2020.609948.

72. Wang, Q.; Zhao, D.; Xian, M.; Wang, Z.; Bi, E.; Su, P.; Qian, J.; Ma, X.; Yang, M.; Liu, L.;, et al. MIF as a Biomarker and Therapeutic Target for Overcoming Resistance to Proteasome Inhibitors in Human Myeloma. Blood 2020, 136, 2557–2573, doi:10.1182/blood.2020005795.

73. Okoye, I.; Xu, L.; Motamedi, M.; Parashar, P.; Walker, J.W.; Elahi, S. Galectin-9 Expression Defines Exhausted T Cells and Impaired Cytotoxic NK Cells in Patients with Virus-Associated Solid Tumors. J Immunother Cancer 2020, 8, e001849, doi:10.1136/jitc-2020-001849.

74. Wu, C.; Thalhamer, T.; Franca, R.F.; Xiao, S.; Wang, C.; Hotta, C.; Zhu, C.; Hirashima, M.; Anderson, A.C.; Kuchroo, V.K. Galectin-9-CD44 Interaction Enhances Stability and Function of Adaptive Regulatory T Cells. Immunity 2014, 41, 270–282, doi:10.1016/j.immuni.2014.06.011.

75. Kam, N.-W.; Lau, C.Y.; Lau, J.Y.H.; Dai, X.; Liang, Y.; Lai, S.P.H.; Chung, M.K.Y.; Yu, V.Z.; Qiu, W.; Yang, M.;, et al. Cell-Associated Galectin 9 Interacts with Cytotoxic T Cells Confers Resistance to Tumor Killing in Nasopharyngeal Carcinoma through Autophagy Activation. Cell Mol Immunol 2025, 22, 260–281, doi:10.1038/s41423-024-01253-8.

76. Kwong, D.L.W.; Kam, N.-W.; Tse, W.C.; Lok, S.H.; Tsao, G.; Lee, V.H.-F. The Tim3-Galectin-9 Interactions in the Tumor Microenvironment of Nasopharyngeal Cancer. Journal of Clinical Oncology 2021, 39, 2629–2629, doi:10.1200/JCO.2021.39.15_suppl.2629.

77. Givant-Horwitz, V.; Davidson, B.; Reich, R. Laminin-Induced Signaling in Tumor Cells. Cancer Res 2004, 64, 3572–3579, doi:10.1158/0008-5472.CAN-03-3424.

78. Winkler, J.; Abisoye-Ogunniyan, A.; Metcalf, K.J.; Werb, Z. Concepts of Extracellular Matrix Remodelling in Tumour Progression and Metastasis. Nat Commun 2020, 11, 5120, doi:10.1038/s41467-020-18794-x.

79. Ramovs, V.; te Molder, L.; Sonnenberg, A. The Opposing Roles of Laminin-Binding Integrins in Cancer. Matrix Biology 2017, 57–58, 213–243, doi:10.1016/j.matbio.2016.08.007.

80. Xu, L.; Wang, B.; Wang, C.; Mao, N.; Huang, Y.; Fu, X.; Feng, T.; He, Q.; Zhang, Y.; You, G.;, et al. A Model of Basement Membrane-Related Regulators for Prediction of Prognoses in Esophageal Cancer and Verification in Vitro. BMC Cancer 2025, 25, 696, doi:10.1186/s12885-025-14081-4.

81. Hu, B.; Ma, Y.; Yang, Y.; Zhang, L.; Han, H.; Chen, J. CD44 Promotes Cell Proliferation in Non-Small Cell Lung Cancer. Oncol Lett 2018, doi:10.3892/ol.2018.8051.

82. Thapa, R.; Wilson, G.D. The Importance of CD44 as a Stem Cell Biomarker and Therapeutic Target in Cancer. Stem Cells Int 2016, 2016, doi:10.1155/2016/2087204.

83. Wu, Z.; Lu, J.; Loo, A.; Ho, N.; Nguyen, D.; Cheng, P.Y.; Mohammed, A.I.; Cirillo, N. Role of CD44 in Chemotherapy Treatment Outcome: A Scoping Review of Clinical Studies. Int J Mol Sci 2024, 25, doi:10.3390/ijms25063141.

84. Feng, E.; Yang, X.; Yang, J.; Qu, Q.; Li, X. LAMB1 Promotes Proliferation and Metastasis in Nasopharyngeal Carcinoma and Shapes the Immune-Suppressive Tumor Microenvironment. Braz J Otorhinolaryngol 2025, 91, 101551, doi:10.1016/j.bjorl.2024.101551.

85. Li, L.; Wei, J.-R.; Dong, J.; Lin, Q.-G.; Tang, H.; Jia, Y.-X.; Tan, W.; Chen, Q.-Y.; Zeng, T.- T.; Xing, S.;, et al. Laminin Γ2–Mediating T Cell Exclusion Attenuates Response to Anti– PD-1 Therapy. Sci Adv 2021, 7, doi:10.1126/sciadv.abc8346.

86. Zhang, P.; Yue, L.; Leng, Q.; Chang, C.; Gan, C.; Ye, T.; Cao, D. Targeting FGFR for Cancer Therapy. J Hematol Oncol 2024, 17, 39, doi:10.1186/s13045-024-01558-1.

87. Peng, M.; Deng, J.; Li, X. Clinical Advances and Challenges in Targeting FGF/FGFR Signaling in Lung Cancer. Mol Cancer 2024, 23, 256, doi:10.1186/s12943-024-02167-9.

88. Nguyen, A.L.; Facey, C.O.B.; Boman, B.M. The Complexity and Significance of Fibroblast Growth Factor (FGF) Signaling for FGF-Targeted Cancer Therapies. Cancers (Basel) 2024, 17, 82, doi:10.3390/cancers17010082.

89. Tay, J.K.; Zhu, C.; Shin, J.H.; Zhu, S.X.; Varma, S.; Foley, J.W.; Vennam, S.; Yip, Y.L.; Goh, C.K.; Wang, D.Y.;, et al. The Microdissected Gene Expression Landscape of Nasopharyngeal Cancer Reveals Vulnerabilities in FGF and Noncanonical NF-ΚB Signaling. Sci Adv 2022, 8, doi:10.1126/sciadv.abh2445.

90. Shen, L.; Li, Y.; Zhao, H. Fibroblast Growth Factor Signaling in Macrophage Polarization: Impact on Health and Diseases. Front Immunol 2024, 15, doi:10.3389/fimmu.2024.1390453.

91. Zhang, Y.; Wang, X. Targeting the Wnt/β-Catenin Signaling Pathway in Cancer. J Hematol Oncol 2020, 13, 165, doi:10.1186/s13045-020-00990-3.

92. Zhou, L.-Q.; Shen, J.-X.; Zhou, T.; Li, C.-L.; Hu, Y.; Xiao, H.-J. The Prognostic Significance of β-Catenin Expression in Patients with Nasopharyngeal Carcinoma: A Systematic Review and Meta-Analysis. Front Genet 2022, 13, doi:10.3389/fgene.2022.953739.

93. Ngernsombat, C.; Prattapong, P.; Larbcharoensub, N.; Khotthong, K.; Janvilisri, T. WNT8B as an Independent Prognostic Marker for Nasopharyngeal Carcinoma. Current Oncology 2021, 28, 2529–2539, doi:10.3390/curroncol28040230.

94. Long, A.; Giroux, V.; Whelan, K.A.; Hamilton, K.E.; Tétreault, M.-P.; Tanaka, K.; Lee, J.-S.; Klein-Szanto, A.J.; Nakagawa, H.; Rustgi, A.K. WNT10A Promotes an Invasive and Self-Renewing Phenotype in Esophageal Squamous Cell Carcinoma. Carcinogenesis 2015, 36, 598–606, doi:10.1093/carcin/bgv025.

95. Larsson, P.; Pettersson, D.; Engqvist, H.; Werner Rönnerman, E.; Forssell-Aronsson, E.; Kovács, A.; Karlsson, P.; Helou, K.; Parris, T.Z. Pan-Cancer Analysis of Genomic and Transcriptomic Data Reveals the Prognostic Relevance of Human Proteasome Genes in Different Cancer Types. BMC Cancer 2022, 22, 993, doi:10.1186/s12885-022-10079-4.

96. Tang, Y.; Guo, Y. A Ubiquitin-Proteasome Gene Signature for Predicting Prognosis in Patients With Lung Adenocarcinoma. Front Genet 2022, 13, doi:10.3389/fgene.2022.893511.

97. Zeng, H.; Geng, X.; Wan, H.; Qu, X.; Tang, S.; Zhang, R.; Zhou, M.; Yu, Z.; Pan, J.; Zheng, H.;, et al. A Molecular Signature of the Ubiquitin-Proteasome System for Forecasting Prognosis in Thyroid Carcinoma Patients. J Inflamm Res 2024, *Volume* 17, 10397–10419, doi:10.2147/JIR.S499820.

98. Chen, X.; Ren, C.; Zhou, Z.; Chen, J.; Fan, X.; Li, X.; Chen, J.; Zhu, J. Development of an Ubiquitin-proteasome System Signature for Predicting Prognosis and Providing Therapeutic Guidance for Patients with Triple-negative Breast Cancer. J Gene Med 2024, 26, doi:10.1002/jgm.3584.

99. Ardo Sanjaya Ubiquitin–Proteasome System Dysregulation in Epstein–Barr Virus-Associated Nasopharyngeal Carcinoma: Associated Codes and Analysis 2025.

